# Genetic advantage and equality of opportunity in education: Two definitions and an empirical application

**DOI:** 10.1101/2021.12.14.472565

**Authors:** Rita Dias Pereira

## Abstract

The literature of Equality of Opportunity (EOp) has long acknowledged the existence of ‘talents’, ‘innate ability’ or ‘genetic ability’. Nonetheless, attempts to explicitly incorporate a measure of innate ability in the quantification of EOp have been rare. On the other hand, the literature of social-science genetics has found credible genetic-based components of EOp, without an explicit quantification of overall EOp. In addition, there exists prevalent disagreement within both kinds of literature on whether innate ability should be perceived as a fair or unfair source of advantage. This paper proposes to quantify EOp while explicitly including a genetic-based measure of innate ability. It proposes two formal definitions of EOp that draw on both stances regarding the compensation of innate ability. Novel testable implications are derived. The educational attainment polygenic index is used as a measure of innate ability while correcting for genetic nurture and accounting for the correlation between genes and other circumstances. An empirical application in the US Health and Retirement Study finds that the share of *inequality* of opportunity is 26% under the view that genetic differences are unfair sources of advantage and 21% otherwise. A comparative analysis over cohorts reveals that the trend in EOp depends on the definition adopted; if genetic advantage is a fair source of inequality then EOp has improved; the opposite holds if one considers genetic advantage an unfair source of inequality. These results highlight the importance of accounting for genetic differences in the EOp framework.

## 1. Introduction

High and increasing education premiums are commonly found worldwide (OECD, 2021; Shambaugh et al., 2017). Moreover, existing research has established an important causal link between higher education, earnings, and better health (Tamborini et al., 2015; Heckman et al., 2018; Van Kippersluis et al., 2011). In this sense, education can be an important instrument for upward social mobility (Chetty et al., 2017). Potentially even more important, education is a human right (Universal Declaration of Human Rights, 1948), and it has intrinsic value. It is clear that fair allocation of education is a necessary building block of a fair society. Governments and organizations devote sizeable resources to evaluate and pursue fairness in education^1^, while top universities boast their meritocratic values^2^.

While the equality of educational opportunity is a widely sought after goal in developed countries, its exact meaning is often ill-defined. It is a common view that parental social background, ethnicity or gender (Lefranc et al., 2009; Hufe et al., 2017; Björklund et al., 2012; Ferreira et al., 2008) are unfair sources of inequality. But what about genetic differences? Although differences in ‘talents’ and ‘genetic’ or ‘innate ability’, are widely acknowledged in the literature of Equality of Opportunity (EOp) (e.g., Rawls, 2009; Dworkin, 1981; Roemer and Trannoy, 2015; Lefranc et al., 2009; Hufe and Peichl, 2020; Hufe et al., 2017), an agreement regarding its classification is far from being reached. Many scholars suggest that genetic-based differences are fair sources of advantage (e.g. Rawls, 1971; Nozick, 1974), some propose that those differences should be considered unfair sources of inequality (e.g. Roemer, 2004; Trannoy, 2019) and others argue that those differences are partly fair and partly unfair (e.g. Lefranc et al., 2009; Lee and Seshadri, 2018).

Today, more than ever, the role of genetic differences is becoming particularly salient. Due to the recent advances of the social-science genetics field, it is now possible to construct credible genetic-based predictions for many traits such as educational attainment (e.g., Harden et al., 2020; Belsky et al., 2016, 2018; Papageorge and Thom, 2020; Barth et al., 2020), intelligence (e.g., Allegrini et al., 2019; Piffer, 2015; Krapohl and Plomin, 2016; Luciano et al., 2017), and even income (Kweon et al., 2020). Despite its increasing use in economics and other fields, a consensus on how to treat these measures from a normative point of view does not currently exist. While some authors conclude that inequalities due to genetic differences are inconsistent with equality of opportunity (Kweon et al., 2020; Conley and Fletcher, 2018) others find that genetic influences on social outcomes can be seen as an index of success in achieving EOp (Rimfeld et al., 2018; Lin, 2020).

Up until now, the EOp literature has extensively attempted to quantify EOp shares without a measure of genetic components. At the same time, the social-genetics literature has extensively attempted to quantify genetic components without a measure of overall EOp. This paper wishes to bridge this gap. First, it proposes two formal definitions of EOp in education that explicitly include genetic differences. The first definition considers genetic differences as an unfair source of advantage – much like social background. The second definition considers genetic differences as a fair source of advantage. Second, it derives novel testable implications for both definitions. And third, it provides an empirical application that quantifies the difference in EOp estimates depending on the definition adopted. This conceptual and empirical contribution aims to clarify existing positions within the equality of opportunity in education debate, as well as to provide an operationalization of EOp measurement under both views.

The conceptual portion of this paper heavily relies on, and extends, the work of Lefranc et al. (2009). The first definition is identical to their strong definition of EOp with genetic differences being added as a standard circumstance. The second definition requires the addition of innate ability as a separate, fourth factor in the definition of EOp.^3^ I derive novel testable implications for the second definition and translate the implications derived in Lefranc et al. (2009) to a regression format. *EOp Definition 1* entails that the explained variance of circumstances should be zero in a situation consistent with EOp. *EOp Definition 2* implies that the incremental explained variance of circumstances above the explained variance of innate ability should be zero in a situation congruent with EOp.

Empirically, this paper exploits recently released genetic data to include a novel measure of innate ability for schooling in the EOp framework: the educational attainment polygenic score (EA PGS from here). Briefly, this score aggregates several genetic variants strongly associated with educational attainment (Lee et al., 2018; Okbay et al., 2016; Rietveld et al., 2013). The lead genetic variants used in the construction of the EA PGS were found to be involved in brain development and neuron-to-neuron communication (Lee et al., 2018). Moreover, within-family and mediation studies clearly show that the EA PGS captures cognitive and non-cognitive abilities (Belsky et al., 2018; Lee et al., 2018; Papageorge and Thom, 2020; Mőttus et al., 2017; Belsky et al., 2016) and thus can serve as a useful proxy for innate abilities. Further, the EA PGS has been shown to predict upward social mobility (Belsky et al., 2016, 2018) and educational attainment within families (Domingue et al., 2015), which contests the view that it is a pure reflection of privilege. Currently, polygenic scores explain up to 13% of the variation in educational attainment (Lee et al., 2018).

The EA PGS has a very attractive feature in the context of the equality of opportunity framework; given that this measure is fixed at conception, it cannot be changed by effort or choice, which results in a clear advantage in comparison to traditional IQ or ability measures often used in the literature. Still, it is well-established that the EA PGS is not a pure proxy for innate ability either: part of the effect of the EA PGS is capturing genetic nurture (Koellinger and Harden, 2018; Kong et al., 2018; Cheesman et al., 2020; Selzam et al., 2019). The idea is that parental genes are transmitted into their children’s genotypes, and these same parental genes may partially shape the rearing environment of the child. Ultimately, the EA PGS will capture a direct effect of the child’s own genes on their outcomes, but will also partially reflect the childhood environment shaped by parents. I follow the reasoning of Wu et al. (2020) to estimate the direct effect of the EA PGS and show how to correct the testable implications.

Empirical results obtained in the Health and Retirement Study (HRS) show that ignoring genetic differences between individuals is far from innocuous and that estimates of EOp with respect to years of education change considerably depending on the definition adopted. In particular, a naive estimation (i.e. ignoring genetic differences) places the inequality share on 20.6%, while under *Definition EOp 1* the share of inequality is 26.2%, and under *Definition EOp 2* this share is 20.9%. A cohort analysis reveals that the marked increase in educational attainment for younger cohorts is not accompanied by a unequivocal increase in EOp. While EOp improves with respect to high school completion under both definitions, the same does not hold for years of education or college completion. The trend in EOp for years of education and college completion depends on the definition adopted. Under *Definition EOp 1* there is persistence of inequality of opportunity, yet under *Definition EOp 2* there is decreasing inequality of opportunity, or in other words, an improvement in EOp. The contrast is explained by the increasing role of innate ability as measured by the direct effect of the EA PGS in explaining college completion. Since the college-earnings premium has increased dramatically in the Unites States (Shambaugh et al., 2017), this raises important questions regarding fairness, and highlights the importance of incorporating genetic advantage in the framework of EOp.

The next section proposes and discusses definitions of equality of opportunity. Section 3 derives testable implications. Section 4 discusses the use of the EA PGS as a novel proxy for innate ability. Section 5 describes the data set and the variables used. Section 6 presents the results and section 7 discusses and concludes.

## 2. Defining Equality of Opportunity in Education

### 2.1. Equality of Opportunity: A short review

Roemer‘s (1993; 1998) formulation of Equality of Opportunity (EOp) decomposes outcomes into *effort* and *circumstances*. Circumstances are aspects that are beyond one’s control, and effort refers to aspects that are within one’s control. This formulation implies that an input of the outcome function is either a circumstance or effort. The population is partitioned into a finite set of *types*, with each type defined by a set of circumstances. EOp is achieved when the outcome is a direct result of effort; that is, circumstances should have no effect on the outcome. Roemer (1993; 1998) has acknowledged that effort might be severely constrained by circumstances. He proposed a measure of *accountable* or *relative* effort which differs from *raw* effort. The idea is that the distribution of effort of a given *type* is independent of an individual’s actions, but rather is a characteristic of the *type*. Since EOp requires there to be no effect by circumstances on the outcome, individuals should not be held accountable for being members of a type with a poor distribution of effort. Rather, individuals should be held accountable for the *rank* they occupy in the distribution of their type. The rank measure of effort effectively nullifies the influence of circumstances on outcomes.^4^

Lefranc et al. (2009) expanded the dichotomous approach of effort and circumstances by proposing a framework that explicitly includes luck as a third factor.^5^ The authors advance three definitions of equality of opportunity; the ‘strong’, the ‘weak’ and the ‘very weak’. The remainder of this paper builds on the ‘strong’ definition.^6^ Under the ‘strong’ EOp definition in Lefranc et al. (2009), justice requires the distribution of outcomes to be exactly the same, for a given level of effort, regardless of circumstances. This excludes any direct or indirect effect - through effort or luck - on outcomes. For example, teacher allocation is at least partially random and is therefore luck. However, the distribution of teacher quality is likely to vary between schools, and school allocation is heavily influenced by circumstances. This type of luck is partially unjust under EOp as it is influenced by circumstances. The only ‘fair’ luck under EOp is a type of luck that affects all individuals evenly, regardless of their circumstances. This does not mean that every individual is affected similarly *ex post*, but rather that each individual is subject to the same prospects *ex ante*. Note that there is an important parallel between effort and luck as circumstances are likely to affect both. Justice according to the ‘strong’ EOp definition does not require a null effect of effort or luck on outcomes; it solely requires that neither is a channel for circumstances to affect outcomes. As Roemer (2003) himself acknowledged, the failure to include luck implies that variables that are not circumstances will inevitably be classified as effort. In this sense, the inclusion of luck is an important step in understanding and evaluating fairness of outcomes. Nonetheless, acknowledging the existence of a third factor implies accepting the inconclusiveness of the size of the residual effort and residual luck; due to their dynamic character, measuring one or the other is an incredibly challenging task.

### 2.2. Defining circumstances, effort and luck

Lefranc et al. (2009) refer to luck as *“situations where individual control, choice or moral responsibility bears no responsibility to the occurrence of outcomes”.* The authors distinguish between four types of luck: (i) social background luck – the type of family one is born into; (ii) option luck – chosen lotteries; (iii) brute luck – non-chosen lotteries; and (iv) genetic luck. This definition of luck is by no means straightforward, as it encapsulates notions of both circumstances and *actual* luck.

This paper does not consider circumstances as a form of luck. Instead, it defines **circumstances** as traits or events that an individual cannot possibly affect. Parental income and parental occupation around birth perfectly fit this definition given that individuals cannot possibly affect these traits. Likewise, genetic constituency also fits this definition, as it is something that one cannot possibly change or influence. In addition, all variables which are determined by time of birth are circumstances, including, for example, the country or city where one is born. **Luck** refers to random factors that the individual affects, even if inadvertently. An example is being hit by lightning. One must choose to go for a walk and to choose a certain path that precisely coincides with the lightning strike. Even though one cannot be held accountable for this sort of luck, one’s choices influence the outcome. Both option luck and brute luck fit into this category, and no further distinctions are made between the two in the remainder of this paper; both are simply referred to as luck. Finally, **effort** entails choices that do not involve a random factor. Effort comprises variables such as the hours devoted to study, hours of physical activity, number of hours worked and effort - physical or mental - exerted to concentrate during those hours.

These definitions imply that not all “traits” can be classified into circumstances, effort or luck. In fact, many outcomes are likely a result of a complex process that depend on circumstances, many types of effort, exerted at different times, and countless combinations of different types of luck. While there are different opinions and choices on how to classify circumstances, effort and luck, the above definitions are objective, and circumvent issues concerning the timing of luck and its correlation with circumstances described in Lefranc and Trannoy (2017). All testable implications derived in section 3 follow mathematically from these definitions.

### 2.3. Two Definitions of Equality of Opportunity with innate ability

This paper proposes two EOp definitions that reflect existing stances on the EOp and social-science genetics literature. *EOp Definition 1* considers innate ability as a standard circumstance. This echoes some of the viewpoints in the EOp literature, such as the one of Björklund et al. (2012)^7^ or Trannoy (2019)^8^. This definition also mirrors some of the positions found in the social-science genetics literature, such as the position of Kweon et al. (2020) or Harden (2021), which treat genetic advantage as a circumstance^9^.

*EOp Definition 2* considers innate ability a special circumstance – one that unlike other circumstances is allowed to influence educational outcomes. This definition parallels Rawls’ original formulation, where “talents and abilities” are considered effort (Rawls, 1971). This view is shared by Nozick (1974) which asserts that individuals are entitled to rewards stemming from their personal, natural endowments. In the social-science genetics literature this stance is found in the work of (Rimfeld et al., 2018) or (Lin, 2020), which treat the influence of genetic advantage on social outcomes as a measure of equality of opportunity. Note that this normative viewpoint has a peculiar implication under existing EOp frameworks: a fair allocation of resources implies a situation where individuals reap the benefits, not of their ability, but of their ability ranking within the distribution of abilities of their type. This is a direct implication of the relative conception of effort described in section 2.1. For innate ability, or talent, to be fully compensated, *EOp Definition 2* requires its addition as a fourth, standalone factor.

This paper will not explore the classification of innate ability as luck, this is, that innate ability should be partially compensated. A thorough discussion on why innate ability should be fully compensated - *EOp Definition 2* -, or not at all - *EOp Definition 1* -, can be found in appendix A.

*EOp Definition 1* is the ‘strong’ EOp definition proposed in Lefranc et al. (2009). While the authors consider genetic ability as an unmeasured form of luck^10^, here innate ability is treated as a circumstance. It states that EOp is satisfied if and only if the distribution of outcomes with respect to the same level of effort is similar, regardless of the circumstances:

#### EOp Definition 1

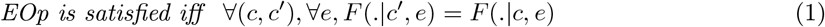

where *c* denotes circumstances, *e* denotes effort and *F* is the outcome function. This definition incorporates luck since otherwise the distribution would collapse to a single mass point.

*EOp Definition 2* adds a fourth factor, innate ability, to the ‘strong’ EOp definition proposed in Lefranc et al. (2009). It states that equality of opportunity is satisfied if and only if, the distribution of educational outcomes is the same for the same level of innate ability and effort, regardless of the circumstances.

#### EOp Definition 2

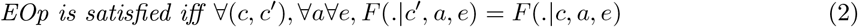

where *a* refers to innate ability.

Figures 1 and 2 depict *EOp Definition 1* and *2* in a Directed Acyclical Graph (DAG). Under *EOp Definition 1*, justice entails that both effort and luck can influence education, and circumstances cannot influence education, neither directly nor indirectly, through effort or luck. Innate ability is included in circumstances. Under *EOp Definition 2*, justice entails that effort, luck and innate ability can influence education and that circumstances cannot influence education, neither directly nor indirectly, through effort or luck. Innate ability is a fourth factor that may influence education, despite it being influenced by circumstances.

**Fig 1.**
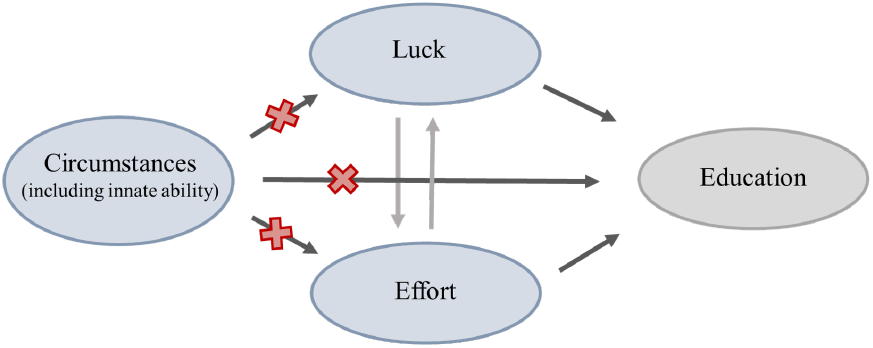
The figure depicts a DAG of a just situation under *EOp Definition 1*.

**Fig 2.**
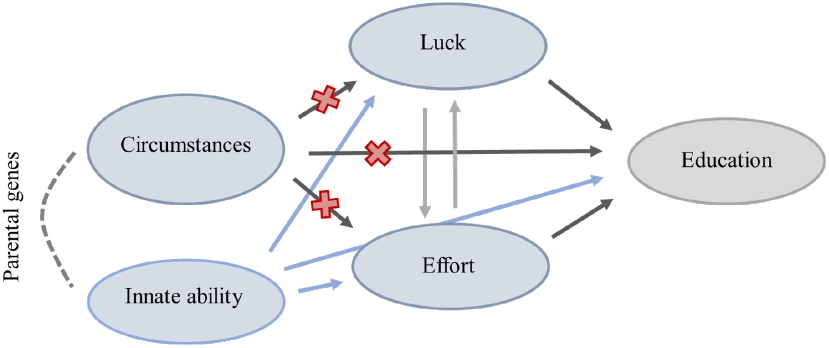
The figure depicts a DAG of a just situation under *EOp Definition 2*.

## 3. Deriving Testable Implications

This section derives empirical testable implications of both *EOp Definition 1* and 2. From here, to avoid confusion, when talking about *equality* of opportunity I will use the acronym EOp, whereas when talking about *inequality* of opportunity I will spell it out.

### 3.1. EOp Definition 1: innate ability as circumstance

*EOp Definition 1* can be simplified if one adheres to the *relative conception of effort*, where the relevant or accountable effort to compensate is the relative effort as measured by the rank measure described in section 2.1. The relative conception of effort is not an assumption but a moral viewpoint; it implies that the relevant effort to be compensated is stripped from the circumstances. A rejection of the conception implies that raw effort is the relevant measure, regardless of the influence of circumstances on effort. Under the relative conception of effort, *EOp Definition 1* can be simplified in the following manner:

#### Result 1.1

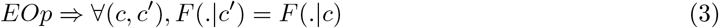

*Result 1.1* entails that adopting the relative conception of effort implies that a measure of effort is no longer required to test EOp. Further, it shows that EOp implies that the distribution of outcomes conditional on circumstances should overlap. The opposite is not true; the overlap of the conditional distributions of outcomes does not imply EOp. The equality on the right is implied by, not equivalent to, EOp. Intuitively, even if the aggregate distributions coincide, luck can be compensated differently depending on circumstances. See Lefranc et al., (2009, p. 1193) for an example of failure of EOp with overlapping conditional distributions. Lefranc et al. (2009) use *Result 1.1* to test EOp by plotting cumulative distributions of income by parental occupational group in France. They find a clear hierarchy between the income distributions by social origin which constitutes a violation of EOp.

While *Result 1.1* is a clever way to test EOp, it also has disadvantages. First, it requires a large sample to correctly plot the conditional cumulative distributions. Second, applying *Result 1.1* when circumstances are continuous is cumbersome since dividing the sample into sub-samples according to types is not straightforward. Finally, comparing EOp across cohorts and countries is not simple. This paper proposes an alternative way to empirically test *EOp Definition 1* that overcomes these three issues.

Consider Decomposition 1, which implies that educational attainment depends on circumstances, effort and luck. The coefficients *α, β* and *θ* depend on the state of the world. The state of the world encapsulates societal and economic variables that are characteristics of a given time period. Societal variables include the degree of gender discrimination or cultural aspects such as knowledge or current beliefs. Economic variables refer to the re-distributive policies, the state of the economy, or incentives put in place by the government. Each cohort is subject to a given state of the world at the same age. Additivity of the independent variables is assumed. This assumption is partially tested in section 6.3. Assume that circumstances are categorical variables varying from 0 to x. An example is parental occupation. This assumption is relaxed later on.

#### Decomposition 1

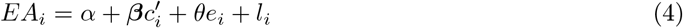

where *EA* is educational attainment of individual *i, c* denotes a vector with measures of circumstances, *e* is the absolute or raw effort, and *l* is the absolute or raw luck. Both absolute effort and luck are expected to be influenced by circumstances. Given that circumstances are by definition predetermined, they must not be influenced by luck or effort. Absolute effort can be re-written as a function of relative effort and the distribution of effort of a given type such that: *e^A^* = *G^t^*(*π_e_*), where *G^t^*(*π_e_*) is the level of effort at the *π_e_* quantile of the distribution of effort of type *t, G^t^*. The same reasoning applies to luck. Absolute luck is a function of relative luck and the distribution of luck of a given type, such that: *l^A^* = *H^t^*(*π_l_*), where *H^t^*(*π_l_*) is the level of luck at the *π_l_* quantile of the distribution of luck of type *t, H^t^*. Recall that the only acceptable sources of inequality are relative effort and relative luck: *π_e_* and *π_l_*. An estimation of Decomposition 1 with unobserved effort and luck yields:

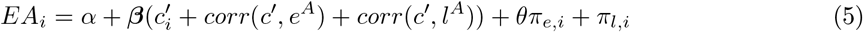

Circumstances *c* shape the distribution of effort - *G^t^* - and luck - *H^t^* - of each type, but not the position of individual *i* in those distributions. In practice this means that corr(*c′, e^A^*) and corr(*c′, l^A^*) are expected to be different from zero while *π_e_* and *π_l_* are independent of circumstances. Given that both effort and luck are unobserved, *β* not only captures the direct effect of circumstances on educational attainment (EA) but also the indirect effect of circumstances through effort *corr*(*c′, e^A^*) and luck *corr*(*c′, l^A^*). The error term will be the sum of the relative effort and luck, *π_e,i_* and *π_l,i_*, which are both considered fair sources of inequality according to *EOp Definition 1*. In this sense, a simple regression of educational attainment on circumstances captures both the i) direct channel of circumstances on EA and ii) indirect channel of circumstances on EA through effort and luck. It’s easy to see that a *β* significantly different from zero constitutes a violation of *EOp Definition 1*. Retrieving the explained variance of equation (5) not only provides a measure of unfair inequality, but at the same time allows cohort and country comparisons. Under *EOp Definition 1*, the variance share of circumstances (*η*) is defined as the *R*-squared of regression 5:

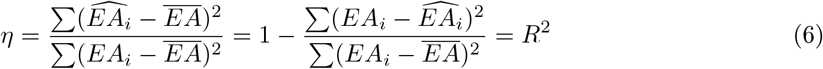

where 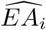 is the predicted educational attainment of individual *i* by the vector of circumstances, 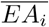 is the average educational attainment for the sample or cohort under analysis and *EA_i_* is the actual educational attainment of individual *i*. By definition, *η* will vary between 0 and 1. *η* will include both the direct effect of circumstances on EA as the indirect effects through effort and luck. An *R*-squared larger than 0 implies that circumstances affect EA, either directly or indirectly, which represents a violation of equality of opportunity according to *EOp Definition 1*. While *η* provides a measure of the share of the outcome that can be explained by circumstances, it cannot distinguish between the direct and indirect effect of circumstances without a measure of effort or luck. Further, *η* can only estimate the size of the two error terms combined; the size of relative effort or relative luck is unknown. The proof that η larger than zero constitutes a violation of *EOp Definition 1* is in Result 1.2:

#### Result 1.2

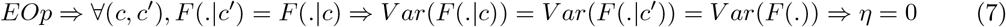

Result 1.2. extends Result 1.1, derived in Lefranc et al. (2009). The result is obtained by placing a variance on both sides of the first equality. The intuition for Result 1.2. is that if the distribution of outcomes is independent of circumstances, the conditional variance should be as well. The derivation is in appendix B. If circumstances are categorical, such as parental occupation, Result 1.2. is weaker than Result 1.1 since it is possible that the explained variance of circumstances is zero, but the conditional cumulative distributions do not overlap. For example, circumstances can explain a zero share of the overall variance of the distribution while explaining other moments of the distribution - for example the skewness or the kurtosis. In that case, plotting the cumulative distribution would reject EOp while relying on *η* would not. This is only true if the circumstances are categorical; this point is elaborated below.

Results 1.1 and 1.2. do not necessarily hold when circumstances are continuous. Plotting the cumulative distributions requires the separation of the sample into sub-samples with different circumstances. It necessarily implies a transformation of continuous circumstances into categorical ones. For example, income in monetary units needs to be divided into high versus low income which implies a loss of information as well as a choice of an arbitrary threshold. A further nuisance is that with multiple circumstances it becomes necessary to create a composite measure of types, based on two or more circumstances (for example, high SES and male, low SES and female). This generates a larger number of types, which compromises the accuracy of the estimation of the cumulative distributions due to sample size reductions. In practice, it becomes unrealistic to test for all possible thresholds. The implication is that cumulative distributions may overlap, yet η may be different from zero. This can be explained by a lack of inequality between types but a presence of inequality within types. For example, the distribution of outcomes between low and high income might overlap yet there might be differences within the “low income” category. In the regression framework there is no need to transform continuous variables in categorical variables and η will capture any differences in conditional means or variances. In sum, if circumstances are continuous, such as income, then the following is true:

#### Result 1.3

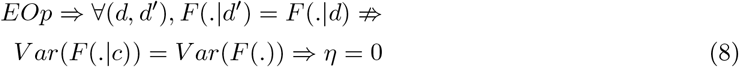

Where *c* refers to a continuous measure of circumstances and *d* refers to a categorical measure of the same circumstance. Briefly, *Result 1.2* holds for categorical circumstances such a parental occupation; overlapping cumulative distributions imply a zero *R*-squared (*η*). *Result 1.3* holds for continuous circumstances that require a transformation for plotting cumulative distributions; overlapping distributions do not imply that the *R*-squared will be zero. This means that, for continuous variables, computing *η* is a stronger test of EOp that plotting cumulative distributions. Note that EOp implies that both hold; the *R*-squared is zero and the cumulative distributions overlap.

The novel testable implications derived in this section are useful as they provide an alternative test that i) does not require the stratification of the population into types; and ii) allows a straightforward comparison of the level of EOp across countries and cohorts. Further, the results derived show that the implications between both tests - plotting cumulative distributions and computing the *R*-squared - vary depending on whether the circumstances are categorical or continuous.

### 3.2. EOp Definition 2: innate ability as special circumstance

This section derives testable implications that parallel the ones in section 3.1, but for *EOp Definition 2*. Under the relative conception of effort, *EOp Definition 2* can be simplified as follows:

#### Result 2.1

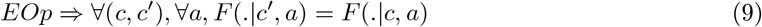

Here, *a* equals innate ability. *Result 2.1.* implies that, conditional on ability, the distribution of outcomes should be the same, regardless of the circumstances. While effort no longer is required for this test, a measure of both circumstances and ability is.

Decomposition 2 entails that educational attainment depends on circumstances, effort, innate ability and luck. Here, a distinction between circumstances and innate ability is made. The coefficients α, *β*, λ and *θ* depend on the state of the world. Additivity of the independent variables is assumed and this is partially tested in section 6.3. Circumstances and innate ability are categorical variables varying from 0 to *x*.

#### Decomposition 2

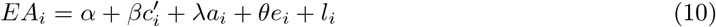

Here, a refers to absolute or raw ability. Absolute effort and luck are expected to be influenced by circumstances and/or innate ability. The opposite is not true, as circumstances and innate ability are predetermined by definition such that they are not influenced by either luck or effort. Using a similar reasoning as in 3.1, estimating *Decomposition 2* with unobserved effort and luck yields:

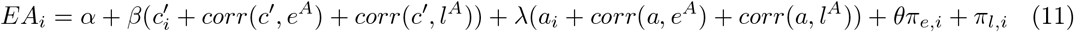

Circumstances *c* and ability a shape the distribution of effort - *G^t^* - and luck - *H^t^* - of each type, but not the position of individual *i* in those distributions.^11^. Both *π_e_* and *π_l_* are independent of circumstances and innate ability by definition. Ability and circumstances are expected to be positively correlated as high ability individuals are expected to acquire a higher social class throughout their life and are more likely to have high ability children. If both circumstances and ability are observed, this is not problematic. Further, from the viewpoint of the individual, both circumstances and ability are determined at the same time which rules out mediation issues. As before, *β* captures the direct and indirect effect of circumstances through effort *corr*(*c′, e^A^*) and luck *corr*(*c^′^, l^A^*). Similarly, λ captures the direct and indirect effect of innate ability on EA. The error term is the sum of the relative effort and luck, *π_e,i_* and *π_l,i_* which are fair sources of inequality according to *EOp Definition 2.* In this case two coefficients are estimated; while *β* captures a violation of EOp, *λ* does not represent a violation of EOp according to Definition 2. The calculation of the variance share of circumstances (*η*) in this case is less straightforward as the two effects need to be disentangled:

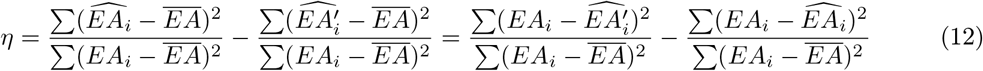

Where 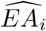 is the predicted educational attainment of individual *i* by the vector of circumstances and innate ability and 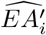 is the predicted educational attainment of individual *i* by innate ability alone. 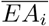 is the average educational attainment for the sample, and *EA_i_* is the actual educational attainment of individual *i*. Since circumstances and innate ability are correlated, under *EOp Definition 2, η* requires a two-step calculation. First, *R*-squared of equation 11 is estimated. Second, the *R*-squared of a regression of educational attainment on innate ability is subtracted. Under justice, innate ability can influence educational attainment directly or indirectly. Circumstances are allowed to influence the outcome, but only through higher innate ability. In this case *η* represents the incremental *R*-squared of circumstances above innate ability. Similarly as in section 3.1, *η* cannot distinguish between the direct and indirect effect of circumstances nor between the the two error terms such that the size of relative effort and relative luck is unknown. The proof that an *η* larger than zero constitutes a violation of *EOp Definition 2* is in Result 2.2:

#### Result 2.2

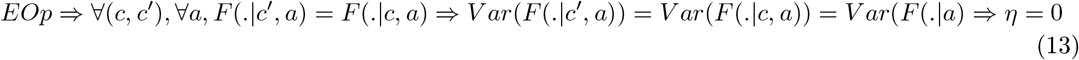

Intuitively, *Result 2.2* entails that, if the conditional distribution of outcomes is independent of circumstances, so should the conditional variance be. The full derivation is in appendix B. It is possible that *η* is zero yet the conditional cumulative distributions do not overlap. The reasoning is the same as the one detailed in subsection 3.1. The difference here is that the sample is split by ability level, such that the cumulative distributions only include individuals with the same ability level. This only holds if circumstances are categorical.

*Results 2.1* and *2.2*. do not necessarily hold when circumstances or innate ability are continuous. Here the assumption that *a* is categorical is dropped. Let’s assume that *a* is a continuous variable that is transformed into a categorical variable for the purpose of plotting cumulative distributions. For example, a continuous measure of innate ability is transformed into quartiles. Figure 3 plots fictitious cumulative distributions of outcomes for two types, for the highest quartile of innate ability.

**Fig 3.**
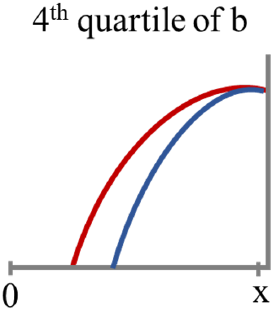
This picture depicts fictitious cumulative distribution functions, for two types. The *x* axis depicts the outcome of interest, that goes from 0 to 1 and the *y* axis depicts the corresponding cumulative distribution. The sample is restricted to people in the highest quartile of ability. The ‘green’ type stochastically dominates the ‘red’ type.

The rank of abilities in picture 3 goes from 75 to 100. Circumstances and ability are expected to be correlated. In that sense, it could be that the average IQ of the green distribution is 81 while the average IQ of the red distribution is 91. This implies that the divergence observed between the cumulative distributions can be entirely driven by ability, rather than by childhood SES. In this case, EOp may hold yet the conditional distributions do not overlap. Under *EOp Definition 2* and continuous ability, it can be misleading to plot cumulative distributions since effectively controlling for the whole spectrum of ability levels is impossible. This is only problematic if ability and SES are correlated. I show that this is indeed the case below.

With respect to circumstances, the reasoning in section 3.1 holds. While plotting the cumulative distributions may show an overlap of the distributions, there could be within-types inequality. Result 2.3. follows:

#### Result 2.3

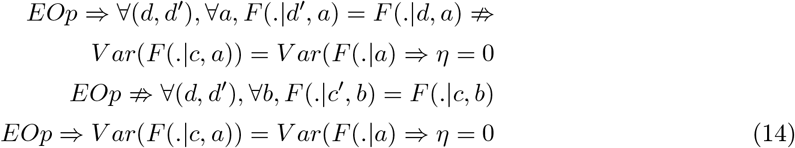

Where *c* refers to a continuous measure of circumstances and *d* refers to a categorical measure of the same circumstance and *a* refers to a continuous measure of innate ability and *b* refers to a categorical measure of the same measure of ability. In short, for categorical circumstances *Result 2.2* holds, and overlapping cumulative distributions imply a null *η*. For continuous circumstances and or ability *Result 2.3* holds; if circumstances are continuous and innate ability categorical then EOp implies *η* = 0 and overlapping cumulative distributions with respect to *d*. However, overlapping distributions do not imply *η* = 0; if circumstances and innate ability are continuous then plotting cumulative distributions is useless and the computation of *η* is the only reliable test of *EOp Definition 2*.

The results derived in this section provide an alternative test for *EOp Definition 2* that does not require the stratification of the population into types. This is crucial for *EOp Definition 2* as plotting conditional cumulative distributions for a continuous measure of ability might lead to misleading conclusions. In particular, one can erroneously reject EOp. The results further propose a measure of EOp that permits a straightforward comparison of the level of EOp across cohorts or countries.

## 4. Proxying innate ability with the educational attainment polygenic score

This section begins by emphasizing that innate ability and later life ability are two distinct concepts. While innate ability can be seen as a latent variable fixed at conception, later life ability is malleable and depends on a complex combination of factors, including personal effort. The distinction between innate and later life ability is found in the work of Trannoy (2019) and Lee and Seshadri (2018). In particular, the papers model ability in time *t* as a function of innate ability, accumulated effort (Trannoy, 2019) and parental and personal investments (Lee and Seshadri, 2018). This reasoning is backed by several findings that suggest that environment can alter gene expression or methylation – the conversion of DNA into proteins (Gibson, 2008; Jaenisch and Bird, 2003; Rakyan et al., 2011) as it can shape gene penetrance – the association between genes and traits without an underlying biological mechanism (Conley et al., 2016). Research also finds that childhood SES predicts brain structure, cognition (Judd et al., 2020), language and executive function (Moorman et al., 2018) independently of genes. In this sense, it very likely that later life ability is a function of effort, circumstances and luck. Accumulated effort such as hours worked, level of focus during those hours, or time spent reading books are examples of how effort can contribute to ability. Poor nutrition, lack of parental nurture, stress, or even negligence are possible pathways through which circumstances may be expected to influence ability. Experiencing an accident that affects brain development or being assigned a good or bad teacher are examples of how luck can affect ability. This highlights that ability in time *t* is expected to be a result of a complex process of initial circumstances, countless random events and accumulated effort during that time. Using a proxy for later life such as IQ measures, grades or achievement tests (Björklund et al., 2012; Plug and Vijverberg, 2003; Von Stumm and Plomin, 2015) implies an inaccurate estimation of EOp, regardless of the definition adopted. If *EOp definition 1* is used, the effort exerted through learning or studying will not be compensated. If *EOp definition 2* is used then the circumstances that contributed to ability formation at a given time will be compensated.

This study proposes to use the educational attainment polygenic score (EA PGS) as a measure of innate ability. Briefly, the EA PGS aggregates several genetic variants that proxy for innate ability and positive personality traits (Smith-Woolley et al., 2019) that predict educational attainment (Okbay et al., 2016; Rietveld et al., 2013; Lee et al., 2018). The lead genetic variants in the score have been shown to be involved in brain development and neuron-to-neuron communication (Lee et al., 2018). Further, this score has been shown to predict educational attainment as well as upward social mobility *within families* (Domingue et al., 2015; Belsky et al., 2016, 2018). Currently, the EA PGS explains up to 13% of the variation in educational attainment (Lee et al., 2018). A brief introduction to human genetics and PGS construction can be found in appendix C.

It is nevertheless well-established that the PGS partially captures environmental effects rather than being a pure biological signal for ability. The idea is that parents both provide genes and influence the environment for a child ; therefore, certain genetic variants of individual x are associated with trait z not because of a direct effect, but rather due to the fact that the mother of individual x has the same genetic variant and therefore provided a certain type of environment that resulted in trait z. Without properly accounting for parental PGS, the child’s PGS will partially capture parental characteristics that shape the rearing environment (Koellinger and Harden, 2018; Kong et al., 2017). This is a standard case of omitted variable bias.

Another potential limitation is that polygenic scores are likely to be under-estimating the true innate ability. In fact, SNP-based heritability estimates suggest that about 22-28% (Davies et al., 2016; Okbay et al., 2016; Tropf et al., 2017) of the variance of educational attainment is explained by additive genetic factors. Hence, current-day polygenic scores explain only about half of this variance. van Kippersluis et al. (2021) show that measurement error of the polygenic score is a key explanation for this difference. It is possible that the predictive power of polygenic scores improves considerably in the future.

Finally, the predictive power of the EA PGS is dependent on social structures; as evidenced by differences in the association between the EA PGS and education for different cohorts (Papageorge and Thom, 2020; Lin, 2020; Herd et al., 2019). This means that while PGSs are “clean” from later life effects that might affect ability but not the genome, the weights attributed to each SNP depend on societal structures that might be discriminatory itself or at least context-dependent. An example could be gender differences: if females are more successful when they are reserved, and males when they are outgoing, the polygenic score will likely miss this heterogeneity and simply yield the average weight of each SNP for men and women. While this is not ideal, it does not imply that PGSs are useless, but rather that they reflect the characteristics that, on average, result in a certain outcome.^12^ The important thing to keep in mind is that the current average depends on societal structures that are malleable, and so the polygenic score should not be interpreted as a fixed, immutable biological characteristic.

It is also important to acknowledge that genetics has a long history of misuse (e.g. Kuhl, 2002). A typical fallacy is that there are undesirable traits. This reasoning is incredibly short-sighted as it disregards the value of diversity, and overlooks talents that might be inherent to those traits wrongly perceived as “inferior”. An example of such a trait is autism, which has been linked with a higher level of attention to detail and a particularly high level of originality in works (Paola et al., 2020). Some companies today have a preference towards autistic individuals, not out of charity or sympathy, but out of an appreciation of their unique attributes (Harris, 2017). This is an important idea, and this work should be read through this lens; while it is clear that some people have a genetic advantage that increases their chances to succeed in the *current* educational system, that does not mean that the current system is not perpetuating a culture of misused potential simply because certain talents are misunderstood or under-valued.

Despite the fact that the research on social science genetics is still developing, the EA PGS is the best, if not only, proxy for innate ability. Since this score is determined at conception, it does not change throughout one’s lifetime.^13^ This effectively ensures that this measure is not contaminated by individual effort nor later life events that are shaped by the individual itself. Nonetheless, it has been shown that this measure partially captures parental genetic endowments. This is not an issue specific to polygenic scores. In fact, relying in an later life measure of ability will suffer from the same issue as parental ability is likely to have shaped the rearing environment of the child and higher ability. The difference is that, while countless factors are likely to influence ability (effort, luck, etc.), only parental genes bias the polygenic score effect size. The reasoning is simple: conditional on the parents, the genes transmitted to a child are completely random, such that it is not possible that effort or luck affect the polygenic score.

By using data sets with adoptees (Cheesman et al., 2020), parental genetic information (Kong et al., 2018) within-family designs (Selzam et al., 2019) offspring phenotypes (Wu et al., 2020), and more recently, within-sibling Genome Wide Analysis (GWAS) (Howe et al., 2021), research estimates that the direct effect is responsible for about 40-50% of the explained variance of the EA PGS on education attainment with the remaining 50-60% being due to genetic nurture. My own estimates from the Health and Retirement Study (HRS) using information on the child’s educational attainment suggest a direct effect of 51.5% (see appendix D for details). To conclude, even though the EA PGS captures environmental pathways, it is possible to have a reliable estimate of the explained variance of the direct effect of the EA PGS given that the parental genome is the only omitted variable.

### 4.1. Including the educational attainment polygenic score in the EOp framework

#### 4.1.1. *EOp Definition 1*: innate ability as a circumstance

Under *EOp Definition 1* innate ability is a standard circumstance and its influence on outcomes is unfair. Using the EA PGS as a proxy for innate ability is therefore rather straightforward. The EA PGS captures both a direct and indirect effect; the direct effect can be perceived as the ‘true’ proxy of innate ability while the indirect effect is capturing parental characteristics that shape the rearing environment. Both the direct and indirect effect are considered circumstances. Given that the interest lies in estimating the total share of variance explained by circumstances, rather than the share of each of the circumstances, the fact that the EA PGS captures both is not an issue. Note that adding the EA PGS as a circumstance implies controlling for a proxy of innate ability as well as for a range of parental genetic-based characteristics, the so-called ‘genetic nurture’. The latter is a novel circumstance in the literature of EOp. Results 1.1 to 1.3 derived in section 3.1 hold when the EA PGS is added as a circumstance.

#### 4.1.2. EOp Definition 2: innate ability as a special circumstance

Under *EOp Definition 2* innate ability is a special circumstance and its influence on outcomes is fair. Including the EA PGS as a proxy for innate ability is slightly more cumbersome. The indirect effect captures parental characteristics that influence the rearing environment. These characteristics do not reflect innate ability *per se* but rather influence later life ability. This paper makes a clear distinction between innate and later life ability, with innate ability being the only circumstance that is permitted a special role (see section 4 for a discussion.) In this sense, only the direct effect of the EA PGS is perceived as the ‘true’ proxy for innate ability and the only component considered a fair source of advantage. Testing EOp under *EOp Definition 2* by plotting conditional cumulative distributions becomes problematic for two reasons. First, an individual level correction of the EA PGS is not yet widely available. This means that conditioning on a certain level of innate ability is not possible. And second, even if it was, the EA PGS is a continuous measure, such that comparing the outcome distributions across circumstance and ability types is not reliable as detailed in section 3.2. Assessing EOp in this case will require the estimation of *η* in a regression framework. In order to exclusively compensate the direct effect of the polygenic score, *η* (equation 12) requires the following adjustment:

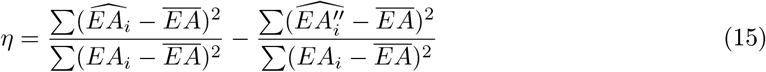

where 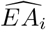 is the predicted educational attainment of individual *i* by the vector of circumstances and EA PGS and 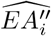 is the predicted educational attainment of individual *i* by the direct effect of the EA PGS alone. 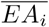 is the average educational attainment. *EA_i_* is the educational attainment of individual *i*. The explained share of circumstances, *η*, can be interpreted as the incremental explained variance of circumstances, over and above the direct effect of the EA PGS. Note that *η* incorporates the indirect effect (or genetic nurture) as a standard circumstance.

Ideally, one would simply have a measure of the direct effect of the polygenic score for each individual. While that might be a reality in the near future, summary statistics of these pioneer studies were not available at the time of writing (Howe et al., 2021). Obtaining an estimate of the direct effect is nevertheless possible but it comes at the expense of additional assumptions. One possible solution is to use the estimates of the direct effect share found in the literature (Selzam et al., 2019; Kong et al., 2018; Cheesman et al., 2020; Wu et al., 2020). These studies generally find that the direct effect accounts for 40-50% of the total explained variance of the EA PGS. Making use of the direct effect share found in the literature relies on the assumption that the direct effect is stable across cohorts and countries and that the direct effect correlates similarly with circumstances, luck and effort (see equation 11 for an intuition). Another potential solution would be to use offspring phenotypic information to estimate the direct and indirect effect of polygenic scores as proposed by Wu et al. (2020). This method assumes that the indirect effect of the polygenic score is stable across generations. While the HRS contains information on the child’s educational attainment, the summary statistics used by Lee and Seshadri (2018) to construct the EA PGS are not publicly available such that a full replication of this method is not possible. Here, *i* follow the reasoning of the authors but rely on an aggregate level correction - at the polygenic score level. The underlying assumption is that this aggregate level correction is a fair approximation for the SNP-level correction suggested by the authors. The details of the construction of the direct effect are in appendix D. The results suggest a direct effect of 51,5%, an estimate in the same ballpark as others found in the literature.

## 5. Data set and variables

### 5.1. Data set

The Health and Retirement Study (HRS) is a longitudinal study that surveys a representative sample of approximately 20,000 people in the United States. It comprises ongoing biennial questionnaires that began in 1992 that focus on understanding work, aging and retirement decisions. This paper uses the “RAND HRS Longitudinal File 2016” which includes questionnaires from 1992 to 2016. The EA PGS was constructed by Lee et al. (2018) and the childhood Socio-Economic Status (SES) measures were constructed by Vable et al. (2017). Both are available on the HRS data products website.

### 5.2. Summary statistics

Columns 1-2 of Table 1 report the summary statistics for the baseline sample. The baseline sample comprises 8,197 individuals of European ancestry^14^ with a non-missing polygenic score who were born between 1920 and 1959 inclusive. The cohort restriction is made to guarantee at least 100 individuals per birth year. Figure A2 in appendix E offers a visual representation of the selection criteria of 100 genotyped individuals by birth year. Cohorts are constructed based on the birth date, with each cohort comprising 10 birth-years. The oldest cohort includes individuals born from 1920 to 1929 and the youngest cohort includes individuals born from 1950 to 1959 inclusive. The cohorts are created with the purpose of increasing sub-sample size and resemble the methods of e.g., Conley et al. (2016), who use cohorts of approximately 2 birth-years, or e.g., Herd et al. (2019), who uses 5 birth-year cohorts. Columns 3-6 of Table 1 report the summary statistics for the oldest and youngest cohort in the baseline sample.

**Table 1.**
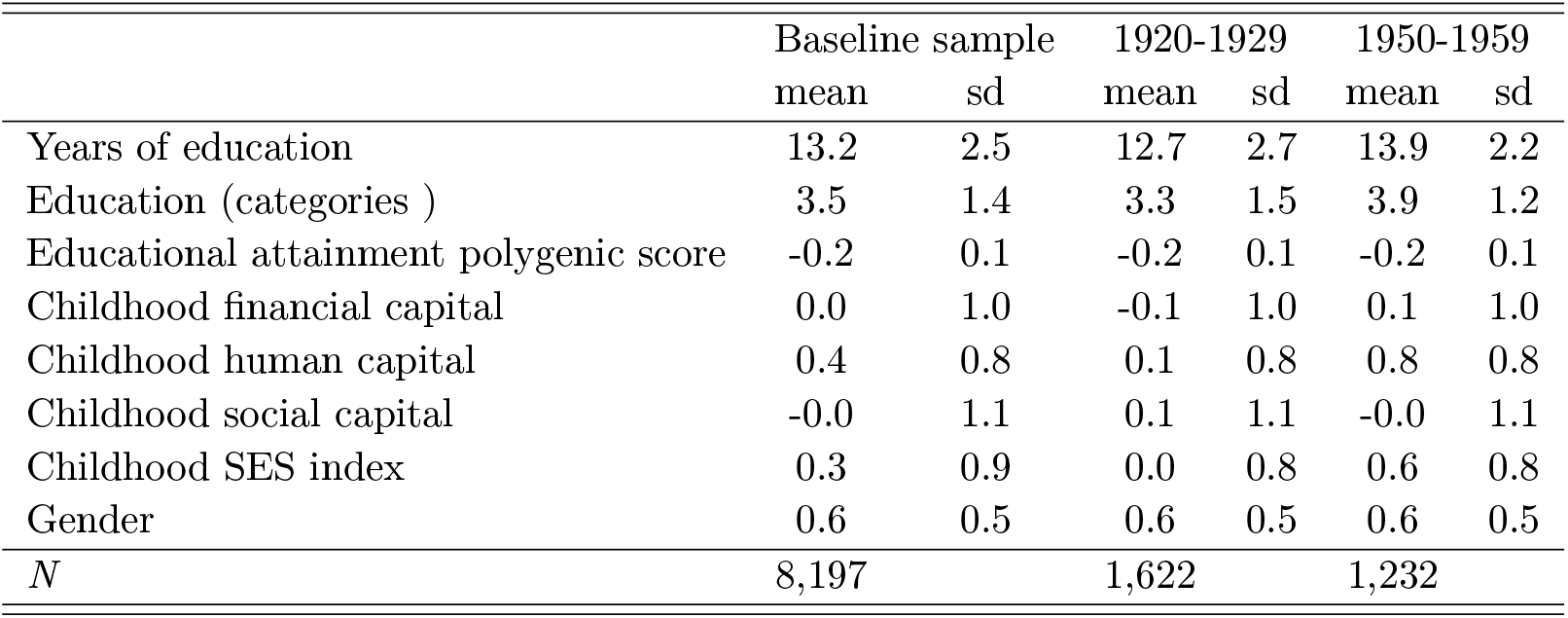
Descriptive statistics of the baseline sample. Columns 1-2 show the baseline sample. Columns 3-4 shows cohorts born between 1926-1930, and columns 5-6 the cohorts born between 1949-1953.

|  | Baseline sample |  | 1920-1929 |  | 1950-1959 |  |
| --- | --- | --- | --- | --- | --- | --- |
|  | mean | sd | mean | sd | mean | sd |
| Years of education | 13.2 | 2.5 | 12.7 | 2.7 | 13.9 | 2.2 |
| Education (categories ) | 3.5 | 1.4 | 3.3 | 1.5 | 3.9 | 1.2 |
| Educational attainment polygenic score | -0.2 | 0.1 | -0.2 | 0.1 | -0.2 | 0.1 |
| Childhood financial capital | 0.0 | 1.0 | -0.1 | 1.0 | 0.1 | 1.0 |
| Childhood human capital | 0.4 | 0.8 | 0.1 | 0.8 | 0.8 | 0.8 |
| Childhood social capital | -0.0 | 1.1 | 0.1 | 1.1 | -0.0 | 1.1 |
| Childhood SES index | 0.3 | 0.9 | 0.0 | 0.8 | 0.6 | 0.8 |
| Gender | 0.6 | 0.5 | 0.6 | 0.5 | 0.6 | 0.5 |
| *N* | 8,197 |  | 1,622 |  | 1,232 |  |

The number of years of education vary between 0 and 17 (not shown). Table 1 shows that the average number of education years is 13.2. The average number of years of education increased from 12.7 to 13.9 years for younger cohorts. Education refers to the level of education, and it varies between 1 and 6. The value 1 corresponds to less than high school, 2 to general education development certificate (GED), 3 to high school graduate, 4 to some college, 5 to a bachelor’s degree and 6 to a master’s degree, a Master of Business Administration (MBA) or a PhD. Figure A7 of appendix E plots the distribution of education levels, for each sub-sample. There is a clear shift of the distribution to the right for younger cohorts, indicating higher levels of education being attained. Childhood financial capital, childhood capital index, childhood social capital and childhood SES index are SES indexes as proposed by Vable et al. (2017). Childhood financial capital is an index that measures average financial resources and financial stability in childhood. Childhood human capital measures maternal and paternal education. Childhood social capital is the sum of family structure (number of household adults) and maternal investment quality (quality of relationship with mother). Average childhood financial capital and human capital increased from −0.1 and 0.1 to 0.1 and 0.8 for younger cohorts and the childhood social capital decreased from 0.1 to 0. This decrease is due to a reduction of maternal investments (from −0.02 to −0.07; not shown). The childhood SES index is the average of the three indexes. All but one component of the childhood SES index improved for younger cohorts such that this index improved for younger cohorts, from 0.0 to 0.6. Finally, gender refers to the proportion of females in the sample. The proportion of females in the baseline sample is 0.6 and this proportion is stable across cohorts.

Figure 4 depicts the aggregate cumulative distribution of education for the youngest – 1920 to 1929 - and oldest - 1950 to 1959 - cohorts in the sample. It is clear from figures 4, A7 and table 1 that education has monotonically increased for younger cohorts, both in years and levels. This is consistent with research that documents an increase in schooling over the same time period (Goldin and Katz, 2008; Lleras-Muney, 2002; Schmidt, 1996; Acemoglu and Angrist, 2000). Unlike educational attainment, the EA PGS has monotonically decreased for younger cohorts as evidenced by figure A4 in appendix E. This finding is consistent with research that finds an increase in educational attainment despite a decrease in the EA PGS (e.g. Courtiol et al., 2016; Beauchamp, 2016). Finally, figure A5 in appendix E depicts a monotonic increase of childhood SES for younger cohorts.

**Fig 4.**
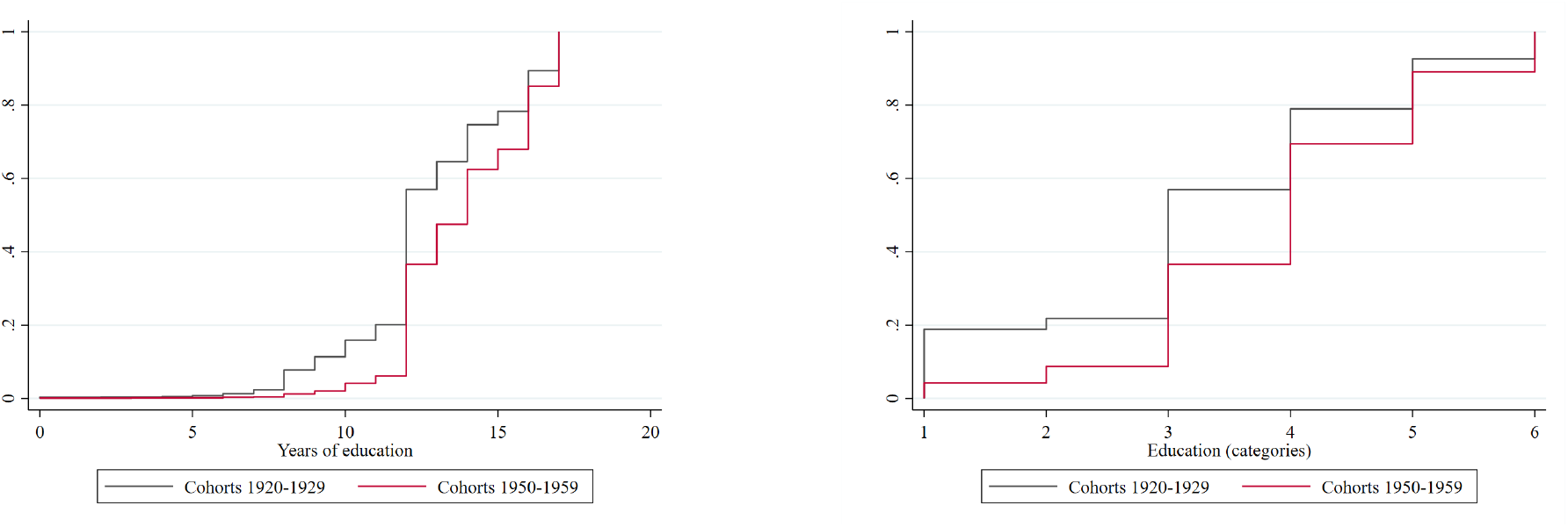
Aggregate cumulative distribution of years and levels of education for the youngest and oldest cohorts.

Figure 5 plots the correlation between childhood SES and EA PGS by cohort. The figure shows a positive and time-varying correlation between the two variables. This positive correlation was at least partially expected. PGS has been shown to predict educational attainment as well as upward social mobility (Domingue et al., 2015; Belsky et al., 2016, 2018). Hence, high EA PGS individuals are more likely to climb the social ladder and, due to biological inheritance, on average have children with a higher PGS. It follows that *on average* the children born in higher SES families have a slightly higher EA PGS. It should be emphasized though that these mean differences across groups are extremely small compared to the within-group differences (see figure A6 in appendix E). Besides, the polygenic score does not exclusively capture cognitive abilities of the child but also the rearing environment provided by the parents. Nonetheless, the positive correlation between SES and the EA PGS is important to our purposes for two reasons. First, it shows that naive estimates of EOp that ignore genetic differences are necessarily incorrect. On the one hand, if genes are considered a circumstance, equality of opportunity estimates are overestimated, as genetic differences will be only partially captured. If, on the other hand, genes are worthy of compensation, then equality of opportunity is underestimated, as differences in outcomes attributed to childhood SES will be partly driven by differential levels of EA PGS. This is contrary to the common claim in the literature that inequality of opportunity estimations always represent a lower bound (Roemer and Trannoy, 2015; García-Gómez et al., 2015; Hufe et al., 2017). The reasoning is simple: an unobserved variable that deserves full compensation which correlates positively with circumstances implies that the role of circumstances is overestimated. Second, the correlation changes over time, which might lead to an incorrect estimate of an EOp trend. An increase in inequality seemingly driven by childhood SES might be caused by the EA PGS alone; either by its increasing correlation with childhood SES, or by its increasing influence on educational attainment or by a combination of both.

**Fig 5.**
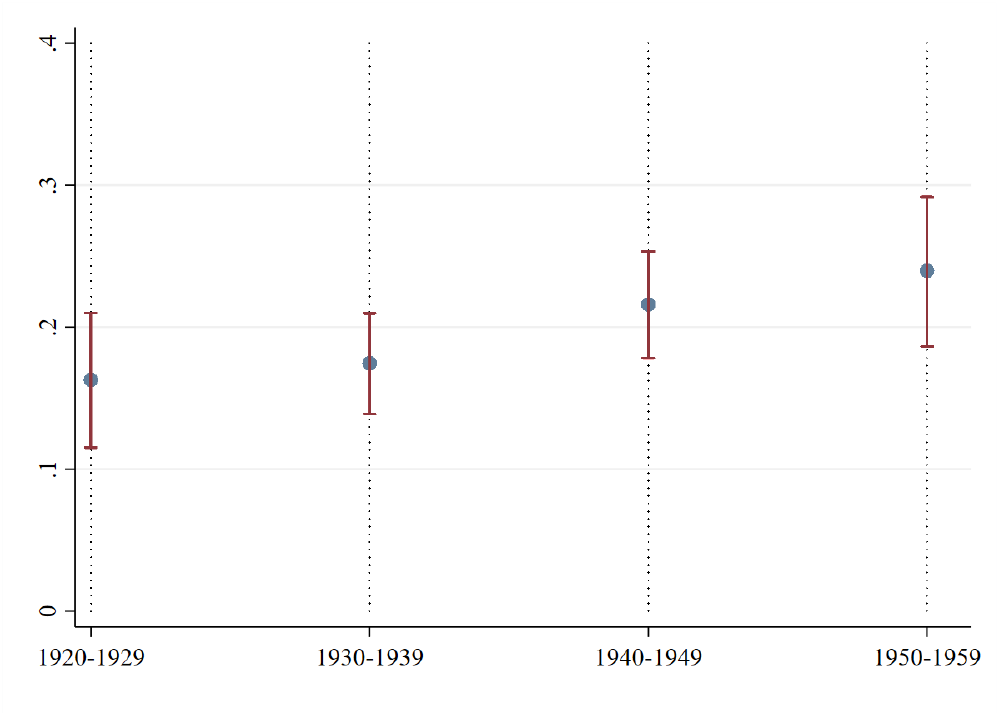
Correlation between the Educational Attainment PGS and childhood SES by cohort.

## 6. Results

This section presents the main empirical results. It computes and compares η, the share of unfair inequality, in educational attainment under *EOp Definition 1* and *2.* It includes a cross-sectional and cohort analysis for both definitions.

### 6.1. Cross-sectional results

Table 2, column 1-3 presents the results of a regression of years of education on childhood SES, gender and on the EA PGS for the baseline sample. A naive estimation of the inequality of opportunity share is 20.6% if genetic differences are ignored (column 1). Under *EOp definition 1*, including genetic advantage as a circumstance increases the share of inequality of opportunity to 26.2%. Under *EOp Definition 2, η* can be calculated with the results of table 2 and the results of appendix D. The direct effect of the EA PGS is considered a fair source of educational inequality such that the explained variance of the direct effect is deducted from the explained variance of all circumstances. *η* with respect to years of education is 20.9 % (0.262 - 0.05315^15^ = 0.209). Hence, ignoring genetic differences results in an underestimation of inequality of opportunity of up to 27%. The inequality share under *EOp definition 1* is 0.053 - the size of the direct effect of the EA PGS - or 25.4% higher than the share estimated under *EOp definition 2*.

**Table 2.** Results of the OLS regression explaining educational attainment. Coefficients are displayed with robust standard errors. Columns 1-3 explain years of education. Columns 4-6 explain high school completion, where 1 is having at least a high school diploma and 0 otherwise. Columns 7-9 explain having a bachelor degree, where 1 is having at least a bachelor degree and 0 otherwise.

|  | Years of education |  |  | High school |  |  | College |  |  |
| --- | --- | --- | --- | --- | --- | --- | --- | --- | --- |
|  | (1) | (2) | (3) | (4) | (5) | (6) | (7) | (8) | (9) |
| Female | -0.29**<br>(0.05) |  | -0.27**<br>(0.05) | 0.02*<br>(0.01) |  | 0.02**<br>(0.01) | -0.10**<br>(0.01) |  | -0.09**<br>(0.01) |
| Childhood financial capital | 0.20**<br>(0.03) |  | 0.17**<br>(0.03) | 0.03**<br>(0.00) |  | 0.03**<br>(0.00) | 0.03**<br>(0.00) |  | 0.02**<br>(0.00) |
| Childhood social capital | 0.06*<br>(0.02) |  | 0.04*<br>(0.02) | 0.01**<br>(0.00) |  | 0.01**<br>(0.00) | 0.01*<br>(0.00) |  | 0.01+<br>(0.00) |
| Childhood human capital | 1.24**<br>(0.03) |  | 1.11**<br>(0.03) | 0.13**<br>(0.01) |  | 0.12**<br>(0.01) | 0.16**<br>(0.01) |  | 0.14**<br>(0.01) |
| EA PGS (standardized) |  | 0.83**<br>(0.03) | 0.61**<br>(0.02) |  | 0.08**<br>(0.00) | 0.06**<br>(0.00) |  | 0.13**<br>(0.00) | 0.10**<br>(0.00) |
| *R*-squared | 0.206 | 0.105 | 0.262 | 0.111 | 0.050 | 0.137 | 0.132 | 0.087 | 0.184 |
| *N* | 8197 | 8197 | 8197 | 8197 | 8197 | 8197 | 8197 | 8197 | 8197 |

Column 3-6 shows the results of a regression explaining high school completion. A naive estimation yields an *η* of 11.1% while including genetic differences under *EOp definition 1* suggests an *η* of 13.7%, a 23% underestimation of inequality of opportunity. *η* for high school completion is estimated under the assumption that the direct effect share is the same as for years of education. This assumption is a necessary one, as there is only information on years of education of the children of the respondents but not on the highest degree completed. In this sense, the assumption is that, for high school completion, the explained variance of direct share of the EA PGS is 2.6% (0.515*0.05=0.026). The consequent inequality of opportunity share of high school attainment is 11% (0.137-0.026=0.111). Coincidentally, this matches the naive inequality of opportunity share of 11.1%.

Finally, columns 7-9 show the results of the regression explaining college completion. A naive estimation puts *η* on 13.2%, while considering genetic differences increases the estimate to 18.4%, implying an underestimation of inequality of opportunity of 39%. Under the same assumption of similarity of the direct effect share, the *η* for college completion is 13.9% (0.184 - 0.515 × 0.087 = 0.139), slightly larger than the naive estimation of 13.2%, suggesting that the naive estimation slightly underestimates inequality of opportunity. Overall, these results highlight the need to discuss the role of genetic advantage in the educational system; this is fundamental for a correct measurement of inequality of opportunity.

Furthermore, table 2 shows that the EA PGS alone explains 10.5% of the educational attainment in years, 5.0% of high school completion and 8.7% of college completion, suggesting a larger role of genetic advantage for higher levels of education.

Finally, table 2 reveals a larger share of unjust inequality for college completion as compared to high school completion, for both definitions. In other words, high school completion is closer to EOp than college completion. Additionally, the share of inequality with respect to years of education is always larger than the share for high school or college completion. A gender comparison in tables A2 and A3 of appendix F reveals slightly larger inequality of opportunity for women. The bulk of inequality is nevertheless explained by differences in childhood SES and the EA PGS.

### 6.2. Cohort analysis results

Figure 6 depicts an estimation of *η* for years of education across cohorts, for both definitions. The figure suggests that, under *EOp definition 1, inequality* of opportunity increased, from approximately 20.2% to 24.5%, although the confidence intervals overlap. Under *EOp definition 2, inequality* of opportunity has decreased, from approximately 19.5% to 11.7%. The increase is statistically significant at the 5% level. For the youngest cohorts, the difference in *η* is the largest; under *EOp definition 1* the inequality of opportunity share is estimated to be twice as large than under *EOp definition 2* - 24.5% versus 11.7%. The difference is driven by an increase in the direct effect of the EA PGS over cohorts, suggesting that the educational system is increasingly compensating genetic advantage. While there was an unequivocal increase in education for younger cohorts, this is not reflected in an unambiguous increase in EOp. Rather, it seems that the improvement in EOp is dependent on how one classifies genetic advantage in the EOp framework; if the direct effect of the polygenic score - my measure of innate ability - is an unfair source of inequality, the prospects seem to be worsening; the opposite is true if the direct effect is considered a fair source of inequality. The cohort analysis results further demonstrate the need to acknowledge and discuss the role of genetic advantage within the EOp framework.

**Fig 6.**
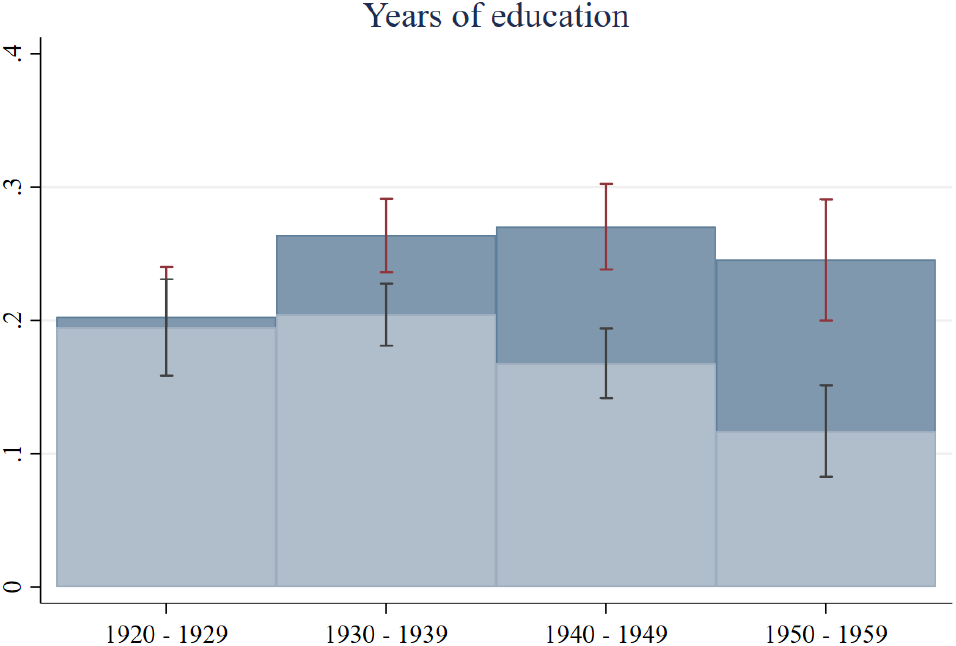
Plot of *η*. The outcome is years of education. In dark blue with dark red I-beams, *η* under *EOp definition 1*; the circumstances included are Childhood SES, EA PGS, and gender. In light blue with grey I-beams, *η* under *EOp definition 2*; the circumstances included are Childhood SES, genetic nurture, and gender. Each bar represents a different cohort. Confidence intervals were constructed using bootstrap with 3000 repetitions.

Figure 7 depicts an estimate of *η* for high school and college completion across cohorts, for both definitions. The figure suggests that *inequality* of opportunity for high school completion has decreased under both definitions although the confidence intervals overlap. The decrease is from 11.9% to 8.7% under *EOp definition 1* and from from 11.4% to 5.3% under *EOp definition 2.* The inequality of opportunity share for college completion increased under *EOp Definition 1* - from 16.4% to 19.8% - and decreased under *EOp Definition 2* - from 15.9% to 7.7%. The decrease under *EOp Definition 2* is statistically significant. Figure 7 suggests that the difference between the trends in inequality of opportunity seen in figure 6 is driven by an increased compensation of genetic advantage at higher levels of education. Overall, the spectacular increase in educational attainment is accompanied by an improvement in EOp in high school completion. However, this does not imply that EOp as a whole has improved; the trend in EOp for college completion and overall years of education is dependent on how one classifies innate ability in the framework of EOp. In fact, it seems that genetic advantage is been increasingly rewarded in higher education.

**Fig 7.**
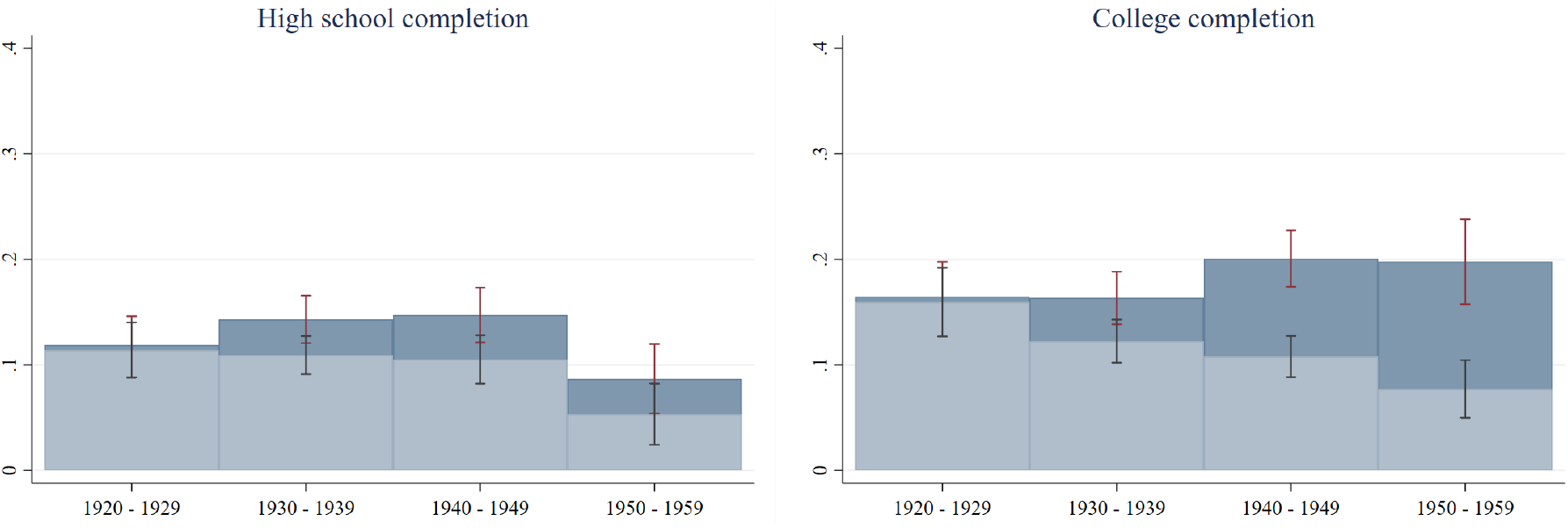
Plots *η.* On the left, high school is the outcome and, on the right, college completion. In dark blue with dark red I-beams, *η* under *EOp definition 1*; the circumstances included are Childhood SES, EA PGS, and gender. In light blue with grey I-beams, *η* under *EOp definition 2*; the circumstances included are Childhood SES, genetic nurture, and gender. Each bar represents a different cohort. Confidence intervals were constructed using bootstrap with 3000 repetitions.

Finally, appendix G plots a naive estimates of *η* over cohorts for years of education, high school and college completion. It is evident that, for every outcome, there are cohorts for which the naive estimation of inequality is considerably higher that estimates under *EOp Definition 2.* This is an unprecedented result in the literature. It is common in the EOp literature to refer to the estimates of inequality of opportunity as lower bounds. This example shows how including a variable that is correlated with circumstances, but that should be compensated, can actually decrease the estimate of inequality. This result further reinforces the importance of including genetic information on estimates of EOp; not only can the estimates be substantially biased, but the bias can go in either direction.

### 6.3. Robustness

Table 3 shows the explained variance of circumstances, η, under different specifications. Columns 1 and 4 show that *η* is considerably larger if *EOp Definition 1* is adopted, for all specifications. Columns 2 and 3 show that *η* increases for younger cohorts under *EOp Definition 1*, irrespective of the specification adopted. On the other hand, Columns 5 and 6 show that *η* decreases under Definition EOp 2 under all specifications but one.

**Table 3.** Estimates of *η* across several specifications. Columns 1 and 4 show the pooled estimate of *η* for years of education under *EOp Definition 1* and 2, respectively. Columns 2 and 3 and Columns 5 and 6 compare *η* for the oldest and youngest cohort in the sample, under *EOp Definition 1* and 2, respectively. The first line shows the original specification. The second a specification where the general cognition PGS is included rather than the EA PGS. The third specification includes the principal components of the genetic relatedness matrix as a circumstance (Price et al., 2006; Rietveld et al., 2014). The fourth specification weights respondents according to the HRS sample weights, averaged over each individual. The fourth and fifth specifications include the survey weights proposed by Domingue et al. (2017) to correct for differential mortality between the genotyped and non-genotyped sample of the HRS.

|  | *EOp Definition 1* |  |  | *EOp Definition 2* |  |  |
| --- | --- | --- | --- | --- | --- | --- |
|  | Pooled | 1920-1929 | 1950-1959 | Pooled | 1920-1929 | 1950-1959 |
|  | (1) | (2) | (3) | (4) | (5) | (6) |
| Original specification | 0.262 | 0.203 | 0.245 | 0.207 | 0.195 | 0.117 |
| Cognition PGS | 0.224 | 0.153 | 0.207 | 0.203 | 0.152 | 0.158 |
| Principal components | 0.268 | 0.209 | 0.246 | 0.213 | 0.201 | 0.118 |
| HRS survey weights | 0.267 | 0.204 | 0.245 | 0.209 | 0.197 | 0.115 |
| Mortality survey weights | 0.270 | 0.199 | 0.278 | 0.214 | 0.191 | 0.128 |
| Birth year survey weights | 0.265 | 0.197 | 0.270 | 0.210 | 0.190 | 0.122 |

The first alternative specification of table 3 uses the cognition polygenic score instead of the EA PGS. The cognition PGS is constructed using the weights from Davies et al. (2018) and is available on the HRS data products website. This score captures cognitive ability as measured by the performance in several cognitive ability tests. This polygenic score is more resilient to environmental changes such as schooling reforms. However, the explanatory power of this score is much lower than educational attainment. In HRS, the EA PGS explains about 11% of the variation in years of education, and the cognition polygenic score explains less than 4% of variation in years of education. This difference is likely to stem from differences in the discovery sample (100,000 versus 1.1 million for educational attainment). In this sense, changes in the explanatory power of the cognition polygenic score are bounded to the explained variance of the score and therefore are overshadowed by changes in the explanatory power of childhood SES. Another limitation of using this score is that a cognitive measure of the offspring is not available such that relying on the direct effect share estimated for years of education is necessary. Under this specification EOp has worsened under both definitions. Nonetheless it is still true that there are non negligible differences in the inequality of opportunity share depending on how one classifies the direct effect of the cognition pgs. The second alternative specification includes the first 10 principal components of the genetic relatedness matrix of each individual (Price et al., 2006) as a circumstance. Briefly, principal components identify structures in the distribution of genetic variation across geographical location and ethnic background. Intuitively, principal components capture genetic ancestries. Including principal components increases *η* marginally under all specifications. The third alternative specification uses sample weights that account for differential probability of selection and differential non-response in each wave^16^ (Sonnega et al., 2014). The HRS survey weights do not account for differential sample selection of the genotyped and non-genotyped sub-samples. The subsequent two specifications use weights that account for differential mortality of these two sub-samples (Domingue et al., 2015). Tables A4 and A5 perform similar robustness checks to the binary outcomes of high school and college completion. The robustness checks confirm that the share of inequality of opportunity is always higher under *EOp Definition 1* as compared to *EOp Definition 2.* Further, all the specifications confirm a decreasing share of inequality of opportunity for high school completion, under both definitions. Finally, all the specifications confirm the trend of increasing inequality under *EOp Definition 1* and decreasing inequality under *EOp Definition 2* for college completion.

Next, the role of non-additive effects is explored by including interactions terms in the OLS regression explaining years of education, for the baseline sample and each cohort. Table A6 in appendix I shows that the incremental explained variance is very small. The largest change in *R*-squared was of 0.4p.p., from 24.5% to 24.9% for the cohort of 1950-1959. Although this mitigates some concerns of sizeable interaction effects between circumstances and the EA PGS, it is not definite proof. First, there might exist other circumstances, not present in the data set, that interact strongly with EA PGS. Further, given that effort and luck are not measured, it is not possible to test possible interactions between circumstances, effort and luck.

Finally, table A7 shows that the *R*-squared is unaffected by distributional changes of the regressors and the outcome.

## 7. Discussion and conclusion

This paper highlights the lack of consensus on how to treat innate differences present in the literature of EOp as in the literature of social-science genetics. While some perceive innate differences as fair sources of advantage, others perceive them as unfair sources of advantage. *Definition EOp 1* considers innate ability a circumstance whose influence on outcomes is deemed unfair, while *Definition EOp 2* considers innate ability a special circumstance, which – unlike other circumstances – should be allowed to influence educational attainment under EOp. Previous quantitative works in the EOp literature have considered innate ability to be either due to effort, circumstance or luck. Conventional ethical views on EOp argue that effort and luck should be neutralized from the effect of circumstances, such that only *relative* levels of effort and luck are compensated. This means that a framework allowing for the full compensation of genetic advantage has been simply non-existent. This paper proposes a novel framework where ability is treated as a standalone factor, such that the *absolute* level of ability could be compensated depending on one’s moral viewpoints.

Novel testable implications of both definitions are derived and applied to the HRS. Results reveal the importance of integrating genetic differences into the literature of EOp. While a naive estimation yields an inequality share (i.e., the explained variance of circumstances) of 20.6%, *Definition EOp 1* places this value on 26.2%, while *Definition EOp 2* suggests an inequality share of 20.9%. The spectacular increase in educational attainment both in years and levels for younger cohorts is reflected in an improvement of EOp for high school completion, under both definitions. Nonetheless, it is not clear whether EOp has improved for college completion or years of education in general. In particular, the trend of inequality of opportunity depends on the definition adopted, if innate talents are unfair sources of inequality there is a deterioration of EOp, and if innate talents are fair sources of inequality, there is an improvement of EOp. Given that the earnings premium of higher education has steadily increased in the U.S. Shambaugh et al. (2017), this finding suggests that a more educated population is not necessarily one that enjoys higher levels of equality of opportunity. If having a college education is increasingly rewarding, and if obtaining a college degree is highly dependent on innate talents, it seems like a discussion on whether innate talents should be rewarded is indeed crucial. The results for college completion and high school hold for several alternative specifications that utilize the general cognition polygenic score, control for genetic ancestry, use a lower bound of the direct effect of the EA PGS and correct for sampling differences of the HRS as a whole and of the genotyped population of the HRS.

One point to keep in mind is that the predictive power of PGS’s is likely to increase in the near future. It is possible that larger GWAS sample sizes or different methods of constructing a PGS are able to increase the explained variance of PGS (McClellan et al., 2017; Visscher et al., 2017; van Kippersluis et al., 2021). This would impact the measure of EOp in several directions, which depend on a) the definition adopted; and b) how strong the new PGS’s correlate with SES. Under definition 1, EOp would never increase since the explanatory power of one circumstance would be increased. The increase in *R*-squared would be at most as large as the additional explained variance of EA PGS. The larger the correlation between the new score and other circumstances is, the less the *R*-squared would increase. Under definition 2, EOp would never decrease as this measure would be allowed to influence the outcome. The extent of the increase would be determined by the correlation between the new score and childhood SES, the smaller the correlation, the less it would decrease, in the limit, it would not decrease at all. In summary, an increase in the explanatory variable of the EA PGS would likely lead to an decrease in EOp under *Definition EOp 1* and to an increase in EOp under *Definition EOp 2*. This implies that, as the predictive power of polygenic scores improves, the larger the difference between the *η* estimates of the two definitions.

One clearly important missing circumstance is ethnicity. The lack of predictive power of polygenic scores for non-European ancestry sub-populations means that the sample is exclusively white which limits the generalization of the findings. Note that the presence of all of the relevant circumstances is not necessary to reject EOp, as having any predictive power of circumstances on outcomes represents a violation of EOp.

As the salience of genetics increases in the private and public sphere, transparent and impartial conversations about the meaning of merit and the role of genetic advantage must take place. In particular, one must question what this study implies for policy makers. First, this study does not endorse a particular ethical stand in the theory of equality of opportunity. It simply wishes to contribute to this discussion by illustrating and clarifying the role of genetic advantage according to the EOp framework; the author acknowledges that the conceptual results might be discouraging for some and may even bring into question the fairness of merit itself (see e.g. Sandel, 2020). Second, the results do not provide support for personalized education based on genetic endowments. In fact, this idea undermines the principle of equality of opportunity as it disregards the role of effort in determining later-life ability and educational attainment. Rather, it is possible to utilize this framework to evaluate *aggregate* levels and trends in EOp and to link those changes with policy interventions and societal changes. In particular, this study finds that an increase in educational attainment does not necessarily imply an increase in EOp in education. In fact, the trend of EOp is dependent on the classification of innate abilities as fair or unfair sources of inequality. This is particularly true for higher levels of educational attainment, which also enjoy of large earnings premium, highlighting the urgency of this discussion.

## A. Normative discussion of innate ability compensation

At a glance, classifying genetic constituency or innate ability as a circumstance feels natural. In fact, people have no responsibility for them and therefore they fit in the standard definition of circumstances in the literature of EOp. The implication of such classification is that innate talents and abilities should not receive compensation under EOp.

While classifying innate ability as a circumstance is instinctive its implications are less so. There are several arguments that support the view that an educational system that rewards innate ability is not necessarily unfair.

An argument put forward in the literature of EOp is the principle of self-ownership. This principle states that agents are entitled to the full benefit of their natural personal endowments such as intelligence, beauty or strength (Nozick, 1974; Vallentyne, 1997). The idea is that your talents are part of one self and therefore should be rewarded.

A second argument is that high ability individuals are likely to be more apt at performing complex tasks and jobs, thus requiring a specific training that is likely to take longer. This does not necessarily imply that being born with high ability should give a person entitlement to higher earnings – this paper adopts no normative viewpoint on this topic. But if one imagines a scenario where people with higher education only have higher earnings to compensate for their delayed entry in the job market, schooling would simply be an allocation mechanism. From a societal point of view, it might be Pareto efficient that higher ability individuals attain higher educational attainment and are allocated to more complex jobs. In reality, education is often used to acquire or maintain status, higher social class and higher earnings. In this sense, one needs to recognize that the education system is not simply an allocation mechanism. Nevertheless, as a society, one needs to realize what is undesirable. Is it that higher ability individuals study longer, or is it that education warrants benefits that are above what is fair? If the first is undesirable, then educational outcomes should not depend on innate ability. If the latter is the issue, then educational outcomes should depend on innate abilities and our educational system and the advantages it provides should be revised.

Lastly, research has suggested that individuals with higher ability have different task (Riding and Read, 1996) and job (Trank et al., 2002) preferences than those with lower ability. Therefore, it is conceivable that low ability individuals derive less utility from education. If this is the case, equalizing outcomes with respect to innate ability would not only undermine personal choice but also would be Pareto inefficient.

The arguments put forward encapsulate either a full or a zero compensation of innate ability which rules out the classification of innate ability as either effort or luck. If ability is categorized as effort or luck, only a relative measure of ability, one that is sterilized from the effects of circumstances, is allowed to influence educational outcomes. In this case only relative ability would matter for an allocation of educational outcomes, regardless of the level of raw or absolute innate ability. This means that a person who would rank high in ability within the distribution of its type would be given a place in education, even if the raw ability is actually lower than an person with better circumstances. In sum, using a relative measure of ability to allocate education would undermine the principle of self-ownership. Additionally, it could sabotage a pareto-efficient allocation of educational attainment.

## B. Proof of Results

### B.1. Result 1.2

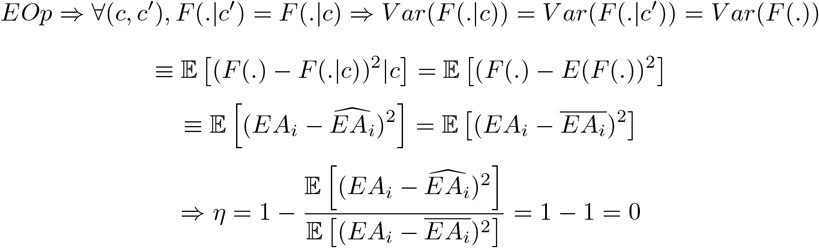

Where *c* refers to the vector of circumstances and *a* to ability. 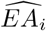 is the predicted educational attainment of individual *i* by the vector of circumstances, 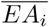 is the average educational attainment for the sample or cohort under analysis and *EA_i_* is the actual educational attainment of individual *i*.

### B.2. Result 2.2

Where *c* refers to the vector of circumstances and a to ability. 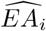 is the predicted educational attainment of individual *i* by the vector of circumstances and the ability and 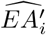 is the predicted educational attainment of individual *i* by the ability alone. 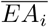 is the average educational attainment for the sample, and *EA_i_* is the actual educational attainment of individual *i*.

## C. Introduction to genetics and polygenic score construction

The human genome is the complete set of genes present in each human cell. It consists of more than 3.2 billion nucleotide pairs located on 23 pairs of chromosomes (Lehrer and Ding, 2017). There are four different base pairs in DNA: adenine, thymine, cytosine and guanine. Approximately 99.6 % of the base pairs are identical for every two unrelated individuals (Kidd et al., 2008). However, some base pairs differ from person to person. The most common genetic variation is called a single nucleotide polymorphism (SNP), and SNPs are the main driver of differences between people. In some rare cases, a difference in a specific locus can single-handedly cause a trait or a disease; Huntington’s disease or Marfan syndrome are examples of such traits. Nonetheless, the majority of traits are polygenic, meaning that they are influenced by many genetic polymorphisms (Chabris et al., 2015). The construction of individual based Polygenic Risk Score (PGS) begins with a genome wide analysis (GWAS) in a discovery sample. A GWAS determines which genetic variants are associated with a certain trait, while controlling for population stratification. Population stratification refers to systematic differences in allele frequencies between sub-populations due to different ancestries. A failure to account for it may lead to false positive or negative associations between the genotype and the trait of interest (Hellwege et al., 2017). Following a GWAS, a PGS can be constructed. The PGS is a sum of SNPs, weighted by the strength of association, and corrected for linkage disequilibrium. Linkage disequilibrium refers to the dependence of gene frequencies at two or more loci where a particular genetic variant is located. It means that SNPs located closer to each other are more likely to be transmitted together (for a more in depth explanation, see e.g. Dudbridge, 2013).

## D. Calculation of the direct effect of the EA PGS

This section estimates the direct effect of the EA PGS by loosely following the reasoning in Wu et al. (2020), which estimates direct and indirect effects using offspring phenotypic information, rather than parental genotypic information. The difference is that, while the authors offer a SNP level correction, I propose a correction at the aggregate level. While a SNP level correction is the most correct one since it corrects the weight of each SNP before the polygenic score construction, the summary statistics used by Lee and Seshadri (2018) are not publicly available which forbids this type of correction. Aggregate corrections are nevertheless often found in the literature (e.g. Cheesman et al., 2020; Muslimova et al., 2020) however, unlike the polygenic scores, which suffer from upward bias, some authors have argued that aggregate corrections suffer from downward bias (see (Trejo and Domingue, 2018)). A recent study that directly measures the direct effect of the EA PGS by conducting a within sibling GWAS has nonethless found a comparable attenuation effect - 46% - as previous estimates in the literature (Howe et al., 2021). This finding partially mitigates the concern that aggregate level corrections suffer from sizeable downward biases.

At the aggregate polygenic score level, the true underlying model is:

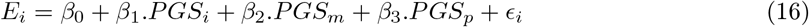

Where *E* refers to the standardized education of the individual *i, PGS_i_* refers to the standardized polygenic score, *PGS_m_* is the maternal polygenic score, *PGS_p_* is the paternal polygenic score and *ϵ_i_* is the error term. Conditional on the paternal and maternal polygenic score, the individual polygenic score is random and therefore *β*_1_, the direct effect, is unbiased. This equation cannot be estimated as the HRS does not have genetic information of trios (mother, father and offspring). Rather, offspring educational attainment is used to estimate the following equations:

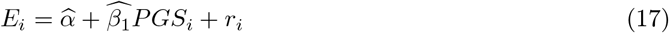

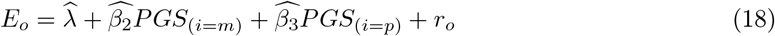

Where *E_o_* is the offspring standardized educational attainment. *PGS*(_*i=m*_) is the individual polygenic score if the individuals is a female and therefore the mother and *PGS*(_*i=p*_) is the individual polygenic score if the individual is male and therefore a father. This estimation is possible since HRS has household level information which makes possible to retrieve couples and their reported offspring educational attainment.

Next, consider that:

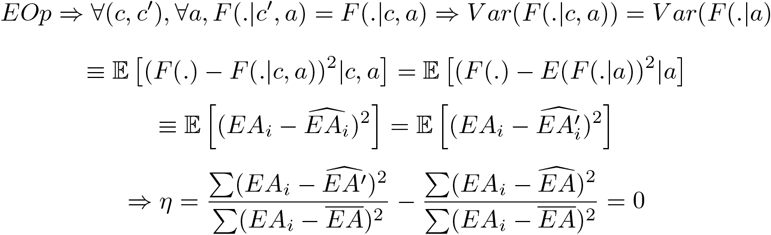

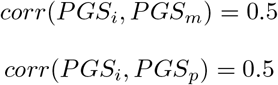

It is expected that the correlation between the individual and its mother or father polygenic score is 0.5 given that half of the parent’s genome is randomly transmitted to the offspring. Empirical estimates confirm this value (e.g. Belsky et al., 2018, find a correlation between the EA PGS of mothers and children of 0.49). Using the HRS I estimate a correlation between the EA PGS of spouses of 0.116 which resembles other estimates found in the literature both in the HRS (e.g. Barban et al., 2016, estimates a correlation of 0.13) as in the UK (e.g. Hugh-Jones et al., 2016, estimates a correlation of 0.11).

The bias of each coefficient can be estimated using the omitted variable formula:

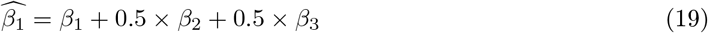

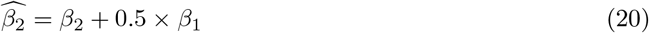

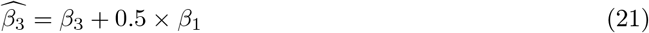

Given that there are three unknowns and three equations, one can retrieve the true betas.

**Table A1.** Results of an OLS regression explaining own and offspring years of education. The first column explains own educational attainment in the baseline sample. The second column explains the average educational attainment of the children of the couple. Mothers and fathers were matched if they a) had the same household identifier, b) the spouse and personal identifier were a match, and c) reported the same educational attainment of their children. Robust standard error in parenthesis.

|  | Years of education<br>(standardized)<br>(1) | Mean offspring years of education<br>(standardized)<br>(2) |
| --- | --- | --- |
| EA PGS (standardized) | 0.320**<br>(0.010) |  |
| EA PGS maternal (standardized) |  | 0.219**<br>(0.021) |
| EA PGS paternal (standardized) |  | 0.255**<br>(0.022) |
| *R*-squared | 0.103 | 0.124 |
| *N* | 8,634 | 1,801 |

Table A1 shows the estimates of 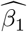 and 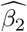 and 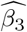, respectively.

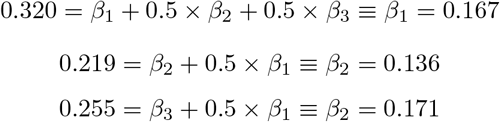

The true *β*_1_ is 0.167, the true *β*_2_ is 0.136 and the true *β_3_* is 0.171. The explained *r*-squared of regression 15 can be decomposed in the following manner:

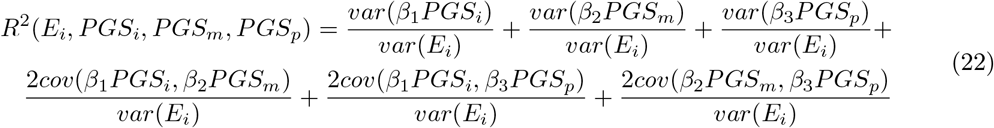

The outcome and regressors are standardized such that *var*(*E_i_*) = 1 and *var*(*PGS_x_*) = 1. If we choose to split the *R*-squared attributed to the covariance equally between its components, the *R*-squared attributed to the direct effect of the polygenic score can be obtained as:

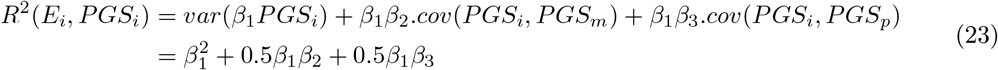

Using the estimated coefficients above, one can estimate the direct effect to be:

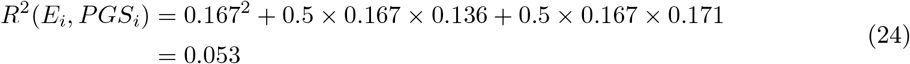

This implies that the share of the explained variance that can be attributed to the direct effect is 0.053/0.103 = 0.515. 0.053 refers to the explained variance of the direct effect of the individual polygenic score, while 0.103 refers to the explained variance of the polygenic score (direct and genetic nurture) in this sample (table A1, Column 1).

Figure A1 depicts 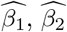 and 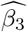 over cohorts. The figure shows an increase of 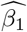 accompanied by a decrease of 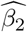 and 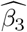, which suggests an increase of the importance of the direct effect of the EA PGS for younger cohorts. Following the reasoning above, one can calculate the explained variance of the direct effect for each cohort. The *R*-squared goes from 0.008 in the cohorts of 1920-1929 to 0.059 in the cohorts of 1930-1939 to 0.102 in the cohorts of 1940-1949 to 0.128 in the cohorts of 1950-1959. This implies that the direct effect share increases from about 8 to 100% over the cohorts. Similar increases were found under different specifications such as a) under 5 year-cohorts division, b) under allocation of families to cohorts exclusively based on the mother or father birth year in order to estimate 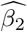 and 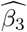, and c) estimation of 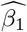 exclusively on the sub sample of couples. Results available upon request. The change of the direct effect over cohorts has not been studied to this date such that a benchmark for comparison does not exist. However, this method did found very similar estimates on average to benchmarks in the literature; it is possible that these averages conceal sizeable differences across cohorts. These stark differences could reveal to be a promising avenue for future research.

**Fig A1.**
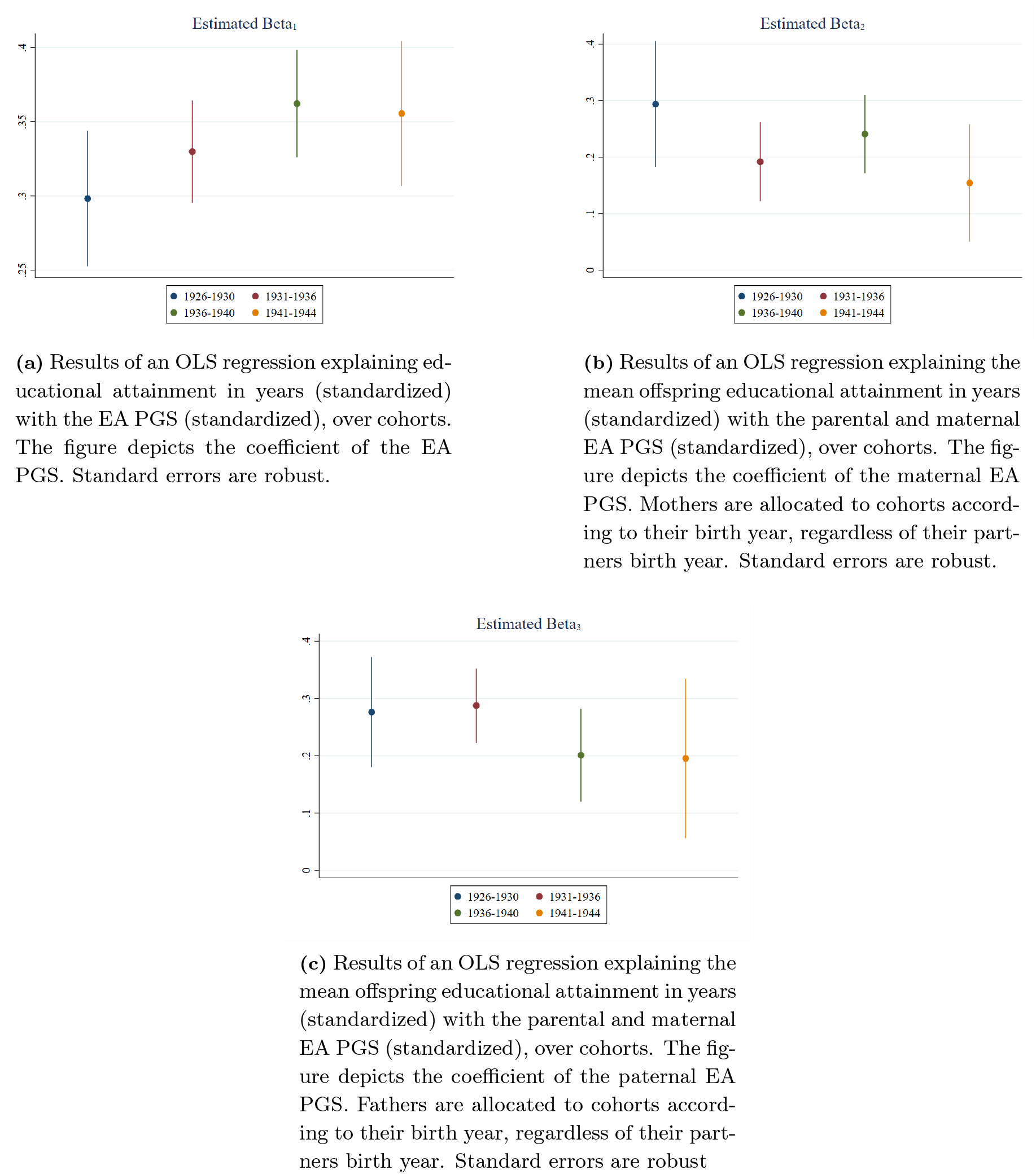
Estimation of 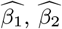 and 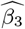 over cohorts

## E. Extended summary statistics

Figure A2 depicts the frequency distribution of the number of observations in the HRS by year of birth, for genotyped and non-genotyped individuals. The distribution by year of birth is similar for the two groups, such that selection by year of birth does not seem to be a concern. A line at 100 observations was added. Only birth years with more than 100 observations were considered for this analysis. This implied including cohorts born between 1920 and 1959 inclusive.

**Fig A2.**
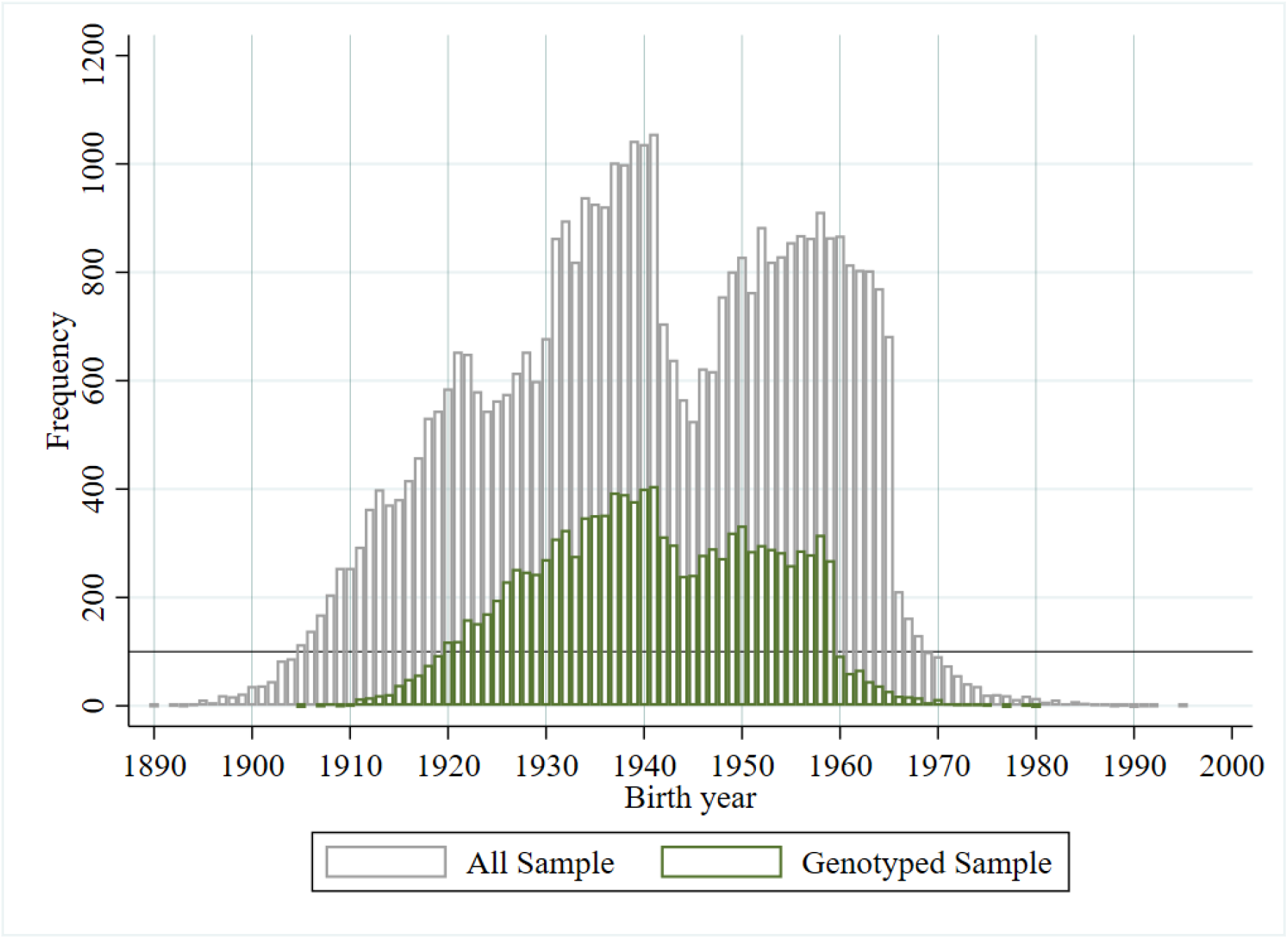
Frequency of observations by year of birth in the HRS.

Figure A7 plots the distribution of education levels for different sub-samples. In the baseline sample, the mode is having at least an high school diploma. There is a clear shift of the distribution to the right for younger cohorts, with a large decrease of people that have less than high school (15p.p. decrease) or just high school (7p.p. decrease) and a large increase of people who have at least some college (11p.p. increase), an increase of individuals who have at least a bachelor’s degree (6p.p. increase) and of individuals who hold a master’s, an MBA or PhD (4p.p. increase). Remarkably, the mode in older cohorts is to be a high school graduate while for younger cohorts is to have some college.

**Fig A3.**
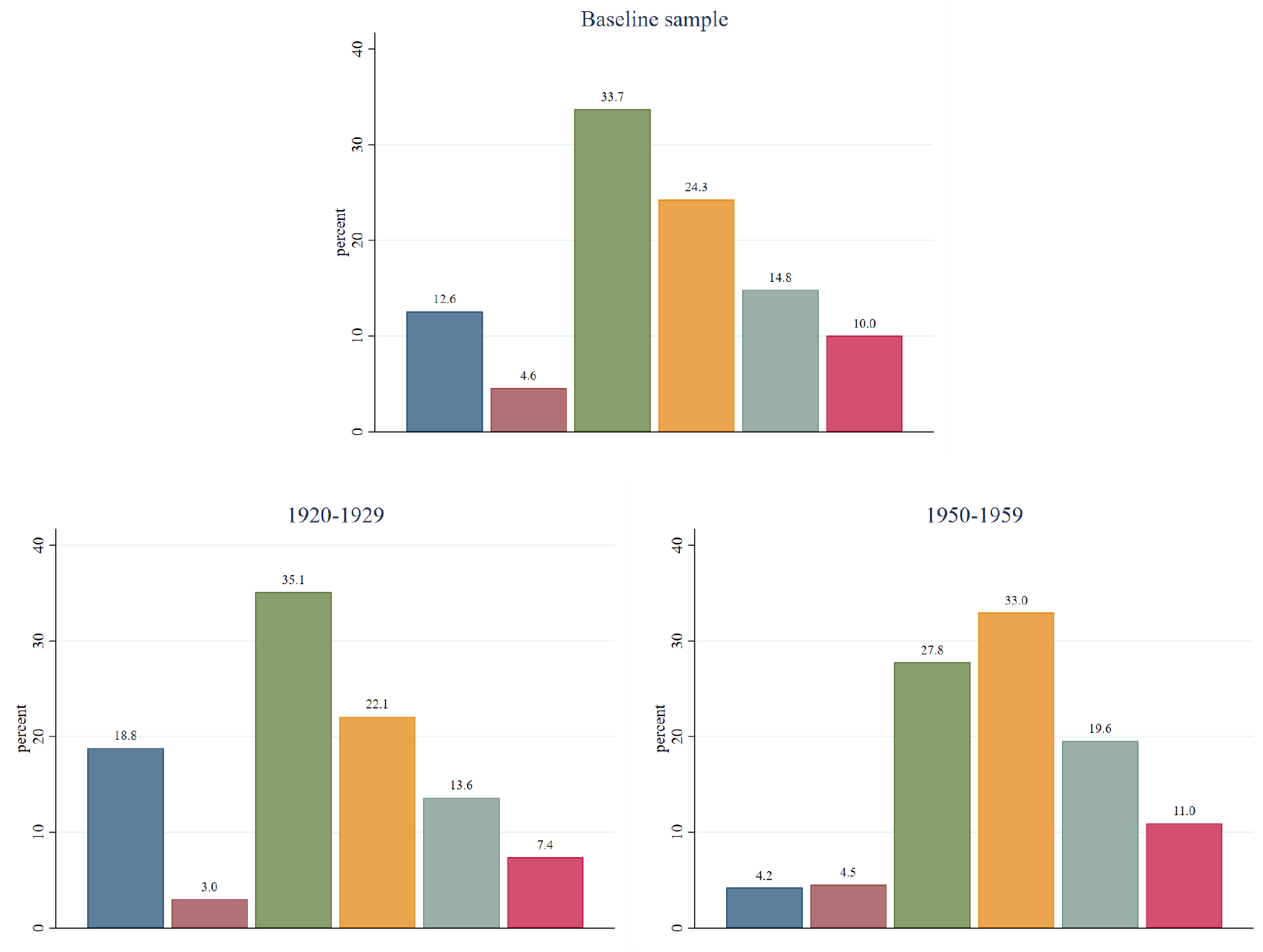
Distribution of education level for different sub-samples. From left to right: less than high school, general education development certificate (GED), high school graduate, some college, bachelor degree and finally master’s degree, a Master of Business Administration (MBA) or phD.

Figure A4 and A5 depict a monotonic decrease of the EA PGS and a monotonic increase of the childhood ses index for younger cohorts. This is evident by the shift of the red cumulative distribution (younger) to the left and right of the blue (older) cumulative distribution, respectively.

**Fig A4.**
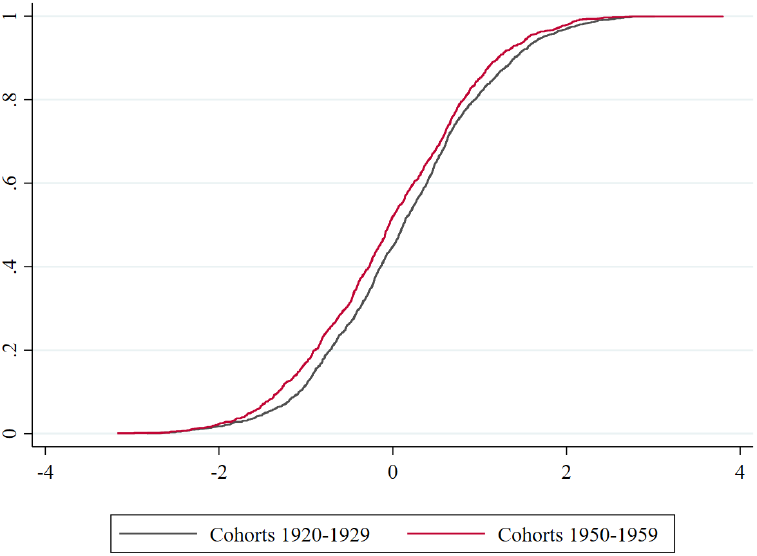
Aggregate cumulative distribution of the EA PGS for the youngest and oldest cohorts.

**Fig A5.**
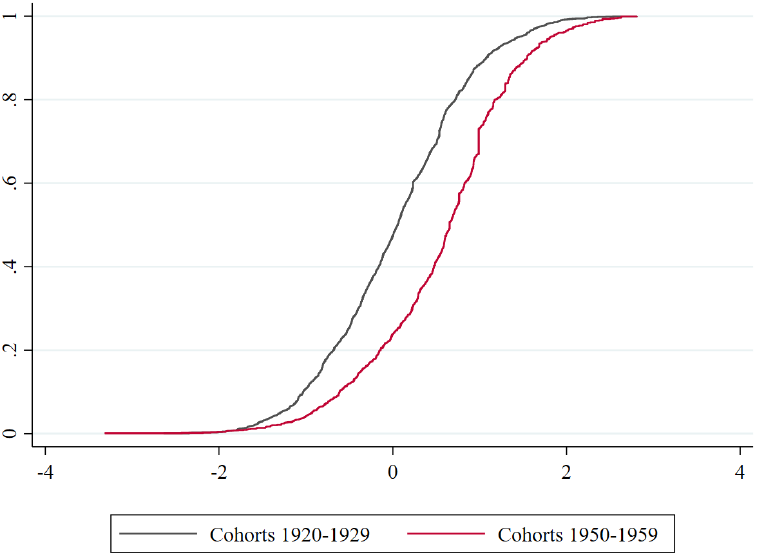
Aggregate cumulative distribution of childhood SES for the youngest and oldest cohorts.

Figure A6 plots the distribution of the EA PGS by the childhood SES. It shows that while a difference in means is observed, a wide dispersion within SES level is obvious.

**Fig A6.**
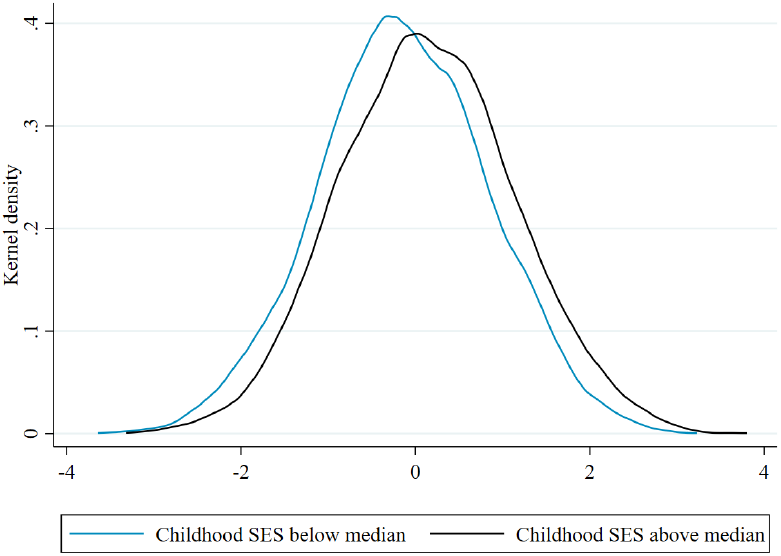
Smoothed distribution of the EA PGS by childhood SES.

## F. Results by gender

Table A2 and A3 show the cross-sectional analysis by gender. It reveals small differences between genders. The largest difference is found on the inequality of opportunity share in years of education; in particular, females are subject to a larger influence of childhood SES on years of education. This can be observed in column 1 and 3 of table A2, where childhood SES explains an additional 4% of variance in years of education for females.

**Table A2.**
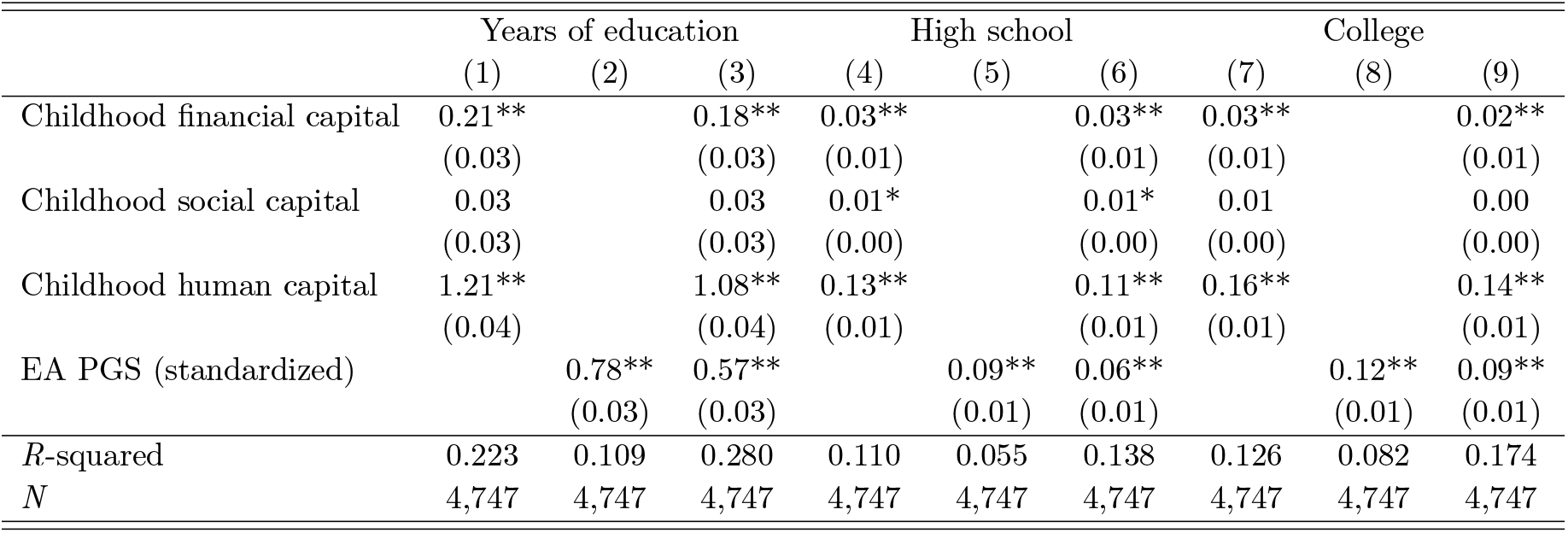
Results of the OLS regression explaining educational attainment of females. Coefficients are displayed with robust standard errors. Columns 1-3 explain years of education. Columns 4-6 explain high school completion, where 1 is having at least a high school diploma and 0 otherwise. Columns 7-9 explain having a bachelor degree, where 1 is having at least a bachelor degree and 0 otherwise.

**Table A3.** Results of the OLS regression explaining educational attainment of males. Coefficients are displayed with robust standard errors. Columns 1-3 explain years of education. Columns 4-6 explain high school completion, where 1 is having at least a high school diploma and 0 otherwise. Columns 7-9 explain having a bachelor degree, where 1 is having at least a bachelor degree and 0 otherwise.

|  | Years of education |  |  | High school |  |  | College |  |  |
| --- | --- | --- | --- | --- | --- | --- | --- | --- | --- |
|  | (1) | (2) | (3) | (4) | (5) | (6) | (7) | (8) | (9) |
| Childhood financial capital | 0.18**<br>(0.04) |  | 0.15**<br>(0.04) | 0.03**<br>(0.01) |  | 0.02**<br>(0.01) | 0.03**<br>(0.01) |  | 0.03**<br>(0.01) |
| Childhood social capital | 0.09*<br>(0.04) |  | 0.07+<br>(0.04) | 0.01*<br>(0.01) |  | 0.01+<br>(0.01) | 0.01+<br>(0.01) |  | 0.01<br>(0.01) |
| Childhood human capital | 1.30**<br>(0.06) |  | 1.16**<br>(0.06) | 0.14**<br>(0.01) |  | 0.13**<br>(0.01) | 0.16**<br>(0.01) |  | 0.14**<br>(0.01) |
| EA PGS (standardized) |  | 0.87**<br>(0.04) | 0.67**<br>(0.04) |  | 0.08**<br>(0.01) | 0.06**<br>(0.01) |  | 0.14**<br>(0.01) | 0.11**<br>(0.01) |
| *R*-squared | 0.183 | 0.101 | 0.240 | 0.114 | 0.046 | 0.136 | 0.113 | 0.093 | 0.172 |
| *N* | 3,450 | 3,450 | 3,450 | 3,450 | 3,450 | 3,450 | 3,450 | 3,450 | 3,450 |

## G. Naive trends of *η*

**Fig A7.**
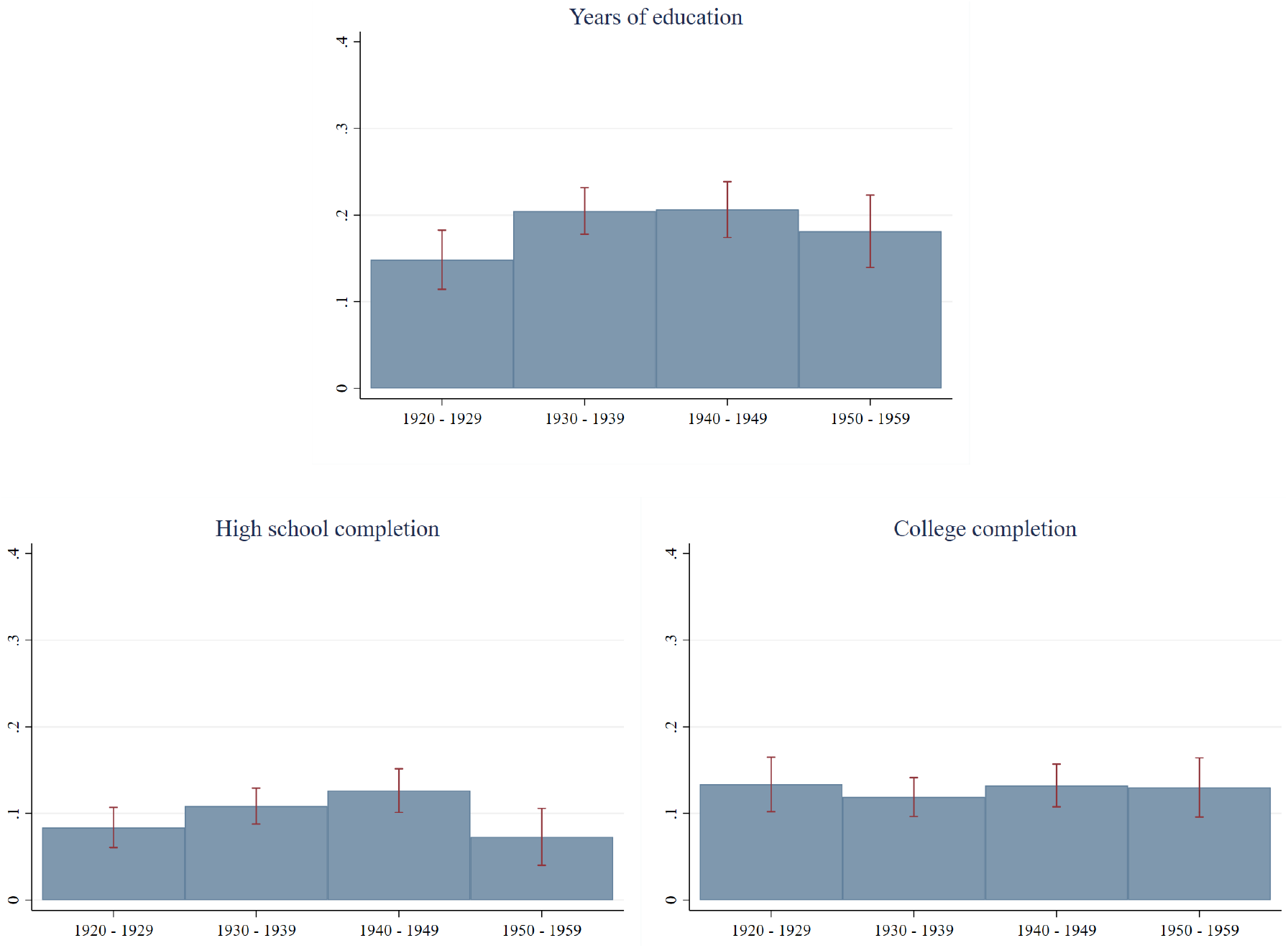
Plots of *η*. On the top, the outcome is years of education. On the left, high school completion and on the right college completion. The circumstances included are Childhood SES, and gender. Each bar represents a different cohort. Confidence intervals were constructed using bootstrap with 3000 repetitions.

## H. Robustness - levels

Tables A4 and A5 depict *η* under several alternative specifications, for high school and college completion, respectively. Columns 1 and 4 show that *η* is larger under *EOp Definition 1* as compared to *EOp Definition 2* for both outcomes. Further, the share of inequality decreased for high school completion under both definitions, for all specifications. The share of inequality in college completion has increased under *EOp Definition 1* and decreased under *EOp Definition 2*. This highlights the indubitable improvement in EOp for high school completion and the importance in classifying innate abilities within the framework of EOp as different classifications yield different conclusions regarding EOp trends. This divergence is explained by the increase in importance of the direct effect for college completion in younger cohorts.

all but one specification: including principal components under *EOp Definition 2*. Overall, this shows that the role of parental SES seems to have increased, but the results are particularly robust to the increase of the role of genetic advantage for college completion.

**Table A4.** Estimates of *η* for high school completion across several alternative specifications. Each specification is detailed in section 6.3. Column 1 and 4 show the pooled estimate of *η* under *EOp Definition 1* and 2, respectively. Column 2 and 3, and 5 and 6 compare *η* for the oldest and youngest cohort in the sample, under EOp *EOp Definition 1* and 2, respectively.

|  | Definition 1 |  |  | Definition 2 |  |  |
| --- | --- | --- | --- | --- | --- | --- |
|  | Pooled | 1926-1930 | 1948-1953 | Pooled | 1926-1930 | 1948-1953 |
|  | (1) | (2) | (3) | (4) | (5) | (6) |
| Original specification | 0.137 | 0.119 | 0.087 | 0.111 | 0.114 | 0.053 |
| Cognition PGS | 0.121 | 0.094 | 0.074 | 0.110 | 0.092 | 0.066 |
| Principal components | 0.145 | 0.127 | 0.091 | 0.119 | 0.122 | 0.057 |
| HRS survey weights | 0.127 | 0.125 | 0.072 | 0.102 | 0.120 | 0.039 |
| Mortality survey weights | 0.148 | 0.124 | 0.099 | 0.120 | 0.119 | 0.056 |
| Birth year survey weights | 0.143 | 0.124 | 0.087 | 0.116 | 0.119 | 0.049 |

**Table A5.** Estimates of *η* for having at least a bachelor degree across several alternative specifications. Each specification is detailed in section 6.3. Column 1 and 4 show the pooled estimate of *η* under *EOp Definition 1* and 2, respectively. Column 2 and 3, and 5 and 6 compare *η* for the oldest and youngest cohort in the sample, under EOp *EOp Definition 1* and 2, respectively.

|  | Definition 1 |  |  | Definition 2 |  |  |
| --- | --- | --- | --- | --- | --- | --- |
|  | Pooled | 1926-1930 | 1948-1953 | Pooled | 1926-1930 | 1948-1953 |
|  | (1) | (2) | (3) | (4) | (5) | (6) |
| Original specification | 0.184 | 0.164 | 0.198 | 0.138 | 0.159 | 0.077 |
| Cognition PGS | 0.148 | 0.138 | 0.152 | 0.133 | 0.137 | 0.112 |
| Principal components | 0.186 | 0.169 | 0.199 | 0.141 | 0.164 | 0.079 |
| HRS survey weights | 0.200 | 0.165 | 0.215 | 0.148 | 0.160 | 0.089 |
| Mortality survey weights | 0.185 | 0.153 | 0.215 | 0.139 | 0.148 | 0.077 |
| Birth year survey weights | 0.185 | 0.150 | 0.214 | 0.140 | 0.145 | 0.079 |

## I. Robustness - interactions

Table A6 compares the *R*-squared of the baseline OLS regressions explaining years of education with the *R*-squared of the OLS regressions with interactions. The largest increase in *R*-squared is for the cohort of 1950-1959, on columns 9 and 10. Including interactions increases the *R*-squared from 24.5% to 24.9%, a 0,4p.p. increase, that corresponds to 1,6% increase.

**Table A6.** Results of an OLS regression explaining years of education. Column 1 and 2 show the results for the whole sample and the subsequent columns show the results for each cohort. Odd columns include childhood SES, gender and the EA PGS as regressors. CFC stands for childhood financial capital, CSC for childhood social capital and CHC for childhood human capital. Pair columns include all interactions as regressors. Robust standard errors in parenthesis.

|  | Baseline |  | 1920-1929 |  | 1930-1939 |  | 1940-1949 |  | 1950-1959 |  |
| --- | --- | --- | --- | --- | --- | --- | --- | --- | --- | --- |
|  | (1) | (2) | (3) | (4) | (5) | (6) | (7) | (8) | (9) | (10) |
| Gender | -0.3**<br>(0.1) | -0.2**<br>(0.1) | -0.2+<br>(0.1) | -0.2<br>(0.1) | -0.3**<br>(0.1) | -0.2*<br>(0.1) | -0.3**<br>(0.1) | -0.3*<br>(0.1) | -0.3*<br>(0.1) | -0.3+<br>(0.2) |
| EA PGS | 0.6**<br>(0.0) | 0.7**<br>(0.0) | 0.7**<br>(0.1) | 0.7**<br>(0.1) | 0.6**<br>(0.0) | 0.7**<br>(0.1) | 0.6**<br>(0.0) | 0.7**<br>(0.1) | 0.6**<br>(0.1) | 0.6**<br>(0.1) |
| CFC | 0.2**<br>(0.0) | 0.2**<br>(0.0) | 0.1*<br>(0.1) | 0.0<br>(0.1) | 0.2**<br>(0.0) | 0.2**<br>(0.1) | 0.2**<br>(0.0) | 0.2*<br>(0.1) | 0.1*<br>(0.1) | 0.2*<br>(0.1) |
| CSC | 0.0*<br>(0.0) | 0.1+<br>(0.0) | 0.0<br>(0.1) | 0.0<br>(0.1) | 0.1*<br>(0.0) | 0.1<br>(0.1) | 0.0<br>(0.0) | 0.1<br>(0.1) | 0.0<br>(0.1) | 0.1<br>(0.1) |
| CHC | 1.1**<br>(0.0) | 1.2**<br>(0.1) | 1.1**<br>(0.1) | 1.1**<br>(0.1) | 1.1**<br>(0.1) | 1.2**<br>(0.1) | 1.1**<br>(0.1) | 1.1**<br>(0.1) | 0.9**<br>(0.1) | 0.9**<br>(0.1) |
| CFC × EA PGS |  | -0.0<br>(0.0) |  | -0.0<br>(0.1) |  | 0.0<br>(0.0) |  | -0.0<br>(0.0) |  | 0.0<br>(0.1) |
| CSC × EA PGS |  | -0.0<br>(0.0) |  | -0.0<br>(0.1) |  | -0.0<br>(0.0) |  | -0.0<br>(0.0) |  | 0.0<br>(0.0) |
| CHC × EA PGS |  | -0.0<br>(0.0) |  | 0.0<br>(0.1) |  | -0.1<br>(0.1) |  | -0.0<br>(0.1) |  | 0.1<br>(0.1) |
| 1.male × EA PGS |  | 0.0<br>(.) |  | 0.0<br>(.) |  | 0.0<br>(.) |  | 0.0<br>(.) |  | 0.0<br>(.) |
| 2.female × EA PGS |  | -0.1+<br>(0.1) |  | -0.2<br>(0.1) |  | -0.1<br>(0.1) |  | -0.0<br>(0.1) |  | -0.2<br>(0.1) |
| 1.male × CFC |  | 0.0<br>(.) |  | 0.0<br>(.) |  | 0.0<br>(.) |  | 0.0<br>(.) |  | 0.0<br>(.) |
| 2.female × CFC |  | 0.0<br>(0.1) |  | 0.2<br>(0.1) |  | 0.0<br>(0.1) |  | -0.0<br>(0.1) |  | -0.1<br>(0.1) |
| 1.male × CSC |  | 0.0<br>(.) |  | 0.0<br>(.) |  | 0.0<br>(.) |  | 0.0<br>(.) |  | 0.0<br>(.) |
| 2.female × CSC |  | -0.0<br>(0.0) |  | 0.0<br>(0.1) |  | -0.0<br>(0.1) |  | -0.1<br>(0.1) |  | -0.0<br>(0.1) |
| 1.male × CHC |  | 0.0<br>(.) |  | 0.0<br>(.) |  | 0.0<br>(.) |  | 0.0<br>(.) |  | 0.0<br>(.) |
| 2.female × CHC |  | -0.1<br>(0.1) |  | -0.0<br>(0.2) |  | -0.2<br>(0.1) |  | -0.1<br>(0.1) |  | 0.1<br>(0.2) |
| Constant | 13.1**<br>(0.1) | 13.1**<br>(0.1) | 12.9**<br>(0.2) | 12.8**<br>(0.2) | 13.0**<br>(0.2) | 13.0**<br>(0.2) | 13.3**<br>(0.2) | 13.3**<br>(0.2) | 13.6**<br>(0.2) | 13.6**<br>(0.3) |
| R-squared | 0.262 | 0.263 | 0.203 | 0.205 | 0.264 | 0.265 | 0.270 | 0.271 | 0.245 | 0.249 |
| N | 8197 | 8197 | 1622 | 1622 | 2880 | 2880 | 2463 | 2463 | 1232 | 1232 |

## J. *R*-squared robustness to distributional changes

Table A7 shows that the *R*-squared is resistant to distributional changes of the regressors and the outcome.

**Table A7.** Results of an OLS regression explaining years of education. In columns 1 and 2 the standardized outcome and in columns 3 and 4 the non-standardized.

|  | Years of education |  |  |  |
| --- | --- | --- | --- | --- |
|  | (non-standardized) | (non-standardized) | (standardized) | (standardized) |
|  | (1) | (2) | (3) | (4) |
| Gender | -0.3**<br>(0.1) | -0.1**<br>(0.0) | -0.3**<br>(0.1) | -0.1**<br>(0.0) |
| EA PGS | 0.6**<br>(0.0) | 0.2**<br>(0.0) |  |  |
| Childhood financial capital | 0.2**<br>(0.0) | 0.1**<br>(0.0) |  |  |
| Childhood social capital | 0.0*<br>(0.0) | 0.0*<br>(0.0) |  |  |
| Childhood human capital | 1.1**<br>(0.0) | 0.4**<br>(0.0) |  |  |
| EA PGS (standardized) |  |  | 0.6**<br>(0.0) | 0.2**<br>(0.0) |
| Childhood financial capital (standardized) |  |  | 0.2**<br>(0.0) | 0.1**<br>(0.0) |
| Childhood social capital (standardized) |  |  | 0.1*<br>(0.0) | 0.0*<br>(0.0) |
| Childhood human capital (standardized) |  |  | 0.9**<br>(0.0) | 0.4**<br>(0.0) |
| *R*-squared | 0.262 | 0.262 | 0.262 | 0.262 |
| *N* | 8,197 | 8,197 | 8,197 | 8,197 |

## Footnotes

1 An example of such is the focus of Organisation for Economic Co-operation and Development (OECD) in pursuing “equal educational opportunities” (Bengtsson, 2008, pg. 1), the European Social Fund, which funds initiatives that promote education of marginalized groups (European Commission, 2018), or the introduction of the Every Student Succeeds Act (2015) in the United States, whose purpose is to provide all children with a significant opportunity to receive a fair and equitable education.

2 For example, Stanford University (2019), University of Oxford (2021) and Cambridge University (2018) have scholarships for lower income students. At the same time, Yale’s president addressed the admission fraud scheme as an “affront to our community’s deeply held values of fairness [and] inclusion” (Salovey, 2019)

3 In EOp, a variable not considered a circumstance is traditionally considered as either effort or luck. However, only relative effort with respect to one’s circumstances ought to be compensated. Something similar happens to luck, as luck is not allowed to be correlated with circumstances. In order to accept the *absolute* level of ability to influence educational outcomes one needs to include it as a fourth factor. A detailed discussion can be found on section 2.3.

4 A technical condition is required for this rank measure of effort to be properly defined. The distribution of absolute effort conditional on circumstances should not exhibit any mass point.

5 While this is far from the first work that proposes luck to be a third factor in the EOp framework (see e.g. Nozick, 1974; Fleurbaey, 1995; Arneson, 1989; Vallentyne, 2002) it is the first one to my knowledge to propose a formal, empirically testable, definition.

6 This choice has several advantages. First, it is the clearest, simplest and most intuitive definition. Second, it simplifies the derivations in section 3.

7 The authors concede that “IQ (…) is most closely tied to merit (in the sense of ability)” (p. 684).

8 Trannoy (2019) distinguishes between current and innate talent. It proposes a cumulative definition of talent where current talent is a function of innate talent and effort. Talent at a given age is partly circumstance, partly effort.

9 While these authors label genetic advantage as 'luck' they effectively treat it as a standard circumstance; its effect on outcomes is labelled as unfair.

10 To be clear, the authors don’t take a normative stance in favour of this classification, rather, they accept that this form of luck can also be considered as a circumstance. However, they do make this choice in their empirical application.

11 innate ability is expected to shape the distribution of luck however it seems less likely that ability influences the distribution of effort. Intuitively, ability is not expected to constrain effort the same way as circumstances do. In particular, circumstances might affect the choice set of effort. One example is a student who needs to take a job after school and therefore has less time to study for his school exam. In this example we can no longer choose between studying 0-10 hours, he can only choose to study 0-3 hours. Ability alone cannot change the choice set of effort. Ability might nonetheless change the productivity of effort, as proposed in Lee and Seshandri (2008). This would imply that *θ* would be ability-dependent. In this sense, *corr*(*a, G^t^*) is expected to be zero while *corr*(*a, H^t^*) is expected to be different than zero.

12 In order to tackle this issue I utilize the cognition PGS in an alternative specification. This PGS is arguably less dependent on societal structures.

13 An exception of a DNA change after conception is called acquired or somatic mutations. These mutations may occur at some time during a person’s life and are present only in specific cells, not in every cell in the body. These mutations can be caused by environmental factors such as ultraviolet radiation from the sun or can occur if an error is made as DNA copies itself during cell division. Mutations acquired in somatic cells (cells other than sperm and egg cells) cannot be passed to the next generation. It is also important to distinguish the difference between DNA and DNA expression. The expression of one’s genome might change throughout one’s lifetime, mostly due to the environment one is exposed to. Nonetheless, one’s genome is unchangeable.

14 Currently, polygenic score prediction in non-European ancestry populations has revealed a low predictive power (Duncan et al., 2019), although efforts are underway to improve upon this. For this study, *i* will focus only on European ancestry individuals.

15 see appendix D for the calculation of the direct effect.

16 The sample weights are meant to account for selection in each wave rather than in each cohort. In particular, the same individuals have several weights, one for each year that he or she answered a questionnaire. To derive individual weights, *i* average the non-missing sample weights for each individual.

